# Sox9-dependent plasticity of esophageal progenitors is fine-tuned by cues from the microenvironment

**DOI:** 10.1101/2025.03.21.644516

**Authors:** Louison Descampe, Benjamin Dassy, Fadi Charara, Ligia Craciun, Laurine Verset, Sheleya Pirard, Quentin Verheye, Morgane Leprovost, Marie Isabelle Garcia, Alizée Vercauteren Drubbel, Benjamin Beck

## Abstract

Cell plasticity governs tissue regeneration but can also drive metaplasia, the replacement of one cell type with another. This process increases the risk of cancer development in several tissues, including esophagus. Esophageal metaplasia development partly depends on keratinocyte plasticity, making regulation of esophageal progenitor fate critical.

We previously identified Sox9 as instrumental in regulating esophageal cell plasticity following Hedgehog pathway activation. Our current study reveals that Hedgehog indirectly regulates Sox9 by modifying epithelial-stromal communication. This activates TGF-β and BMP pathways in epithelial cells, which synergistically regulate Sox9 and stimulate a transcriptomic program resembling squamo-columnar junction progenitors, which are prone to initiate metaplasia. Importantly, we demonstrate pharmacological modulation of this plasticity *in vivo*. Indeed, Ibuprofen inhibits Hedgehog-induced Sox9 expression by directly targeting epithelial cells, providing proof of concept for pharmacological intervention in cell plasticity with implications for regenerative medicine and metaplasia treatment.

## Introduction

Cell plasticity refers to the ability of cells to change their identity or function in response to various stimuli, such as injury or inflammation. In some cases, cell plasticity can lead to metaplasia, which is defined as the replacement of one cell type with another. Metaplasia can be seen as an adaptative process to tissue damage or inflammation, but it can also be a precursor to cancer ^1^. For example, in the esophagus, chronic gastro-esophageal reflux disease (GERD) can trigger the replacement of squamous cells with columnar cells, leading to metaplasia ^2^. This columnar metaplasia is a risk factor for the development of Barrett’s esophagus (BE), a precancerous condition that can lead to esophageal adenocarcinoma.

The finding that p63+ squamous epithelial cells can give rise to intestinal-like metaplasia in a surgical model of GERD ^3^ suggests that keratinocytes can indeed adapt to chronic reflux. Interestingly, squamous progenitors at the squamo-columnar junction (SCJ), characterized by Keratin7 expression, can transdifferentiate upon overexpression of Sox2 or human CDX2 ^3^, implying that they may work in concert with columnar epithelial cells at the SCJ to form columnar metaplasia ^1^. These progenitors seem more competent to initiate metaplasia than the rest of foregut keratinocytes, suggesting that SCJ squamous progenitors are highly plastic. Ablative strategies against metaplasia, such as submucosal endoscopic dissection ^4^, may also impact the fate of esophageal progenitors. Characterization of the mechanisms regulating progenitor plasticity need to be studied in more detail to improve our understanding of esophageal tissue regeneration and repair, particularly the role of the microenvironment.

Previous studies demonstrated that the Hedgehog (HH) pathway, which is turned off in embryonic foregut bipotent progenitors once they differentiate into squamous lineage, is reactivated in esophageal progenitors upon chronic acid reflux in mice and humans ^5,6^. We recently reported that the HH pathway is constitutively activated in SCJ squamous progenitors and that the sole reactivation of this pathway in esophageal cells stimulates cell plasticity by turning on an embryonic-like transcriptomic program before activating differentiation into columnar epithelium ^6^. Notably, we identified the transcription factor Sox9 as playing a crucial role in the regulation of esophageal cell plasticity. Although BMP4 has been proposed to regulate Sox9 expression in esophageal keratinocytes ^5^, the mechanisms underlying Sox9 appearance in esophageal cells *in vivo* remain unknown.

In this study, we discovered that the expression of Sox9 in mouse esophageal progenitors is indirectly regulated by the HH pathway. Through a combination of *in silico* analysis of the open chromatin region in the Sox9 promoter, analysis of single cell RNA sequencing data with a focus on cell-cell interactions, and *in vitro* experiments utilizing esophagus epithelium organoids, we found that TGF-β is a key regulator of Sox9 expression. In our model, BMP alone does not directly promote Sox9 expression but rather reinforce the action of TGF-β on keratinocytes. Transcriptomics data suggest that TGF-β is activated by several mechanisms including modifications of the extracellular matrix, which stimulate the release of latent TGF-β. The sole co-stimulation of esophageal progenitors with BMP and TGF-β *in vitro* is sufficient to turn them into Keratin7+ keratinocytes that resemble HH-stimulated keratinocytes, as well as SCJ squamous progenitors, which have the competence to initiate specialized metaplasia *in vivo*.

Finally, employing a transgenic mouse model, bulk and single-cell RNA-seq, and organoids, we demonstrated the possibility of repressing Sox9 expression in esophageal cells pharmacologically using non-steroidal anti-inflammatory drug both *in vivo* and *in vitro*. Intriguingly, this mechanism operates independently of the TGF-β pathway and mostly through Sox9 protein stabilization in epithelial cells. These findings underscore the intricate crosstalk between epithelial cells and their underlying mesenchyme, elucidating how their interactions impact cell plasticity in the esophagus. Furthermore, they illustrate how pharmacological tools could be employed to maintain keratinocytes in their squamous fate.

## Results

### Sox9 expression in esophageal keratinocytes is indirectly regulated by the Hedgehog pathway

Under physiological conditions, Sox9 is absent from foregut keratinocytes except at the SCJ (Fig. 1a and Supp. Fig. 1d). In the Krt5CreER:RosaSmoM2 (K5:Smo) transgenic mouse model, tamoxifen injection induces HH pathway activation in esophageal basal progenitors (Fig. 1b and Supp. Fig. 1a-c) ^6^. Sox9 is detected in some basal cells 5 days after tamoxifen induction and its expression subsequently further increases in clusters of basal keratinocytes (Fig. 1c and Supp. Fig. 1e-g).

**Fig. 1.**
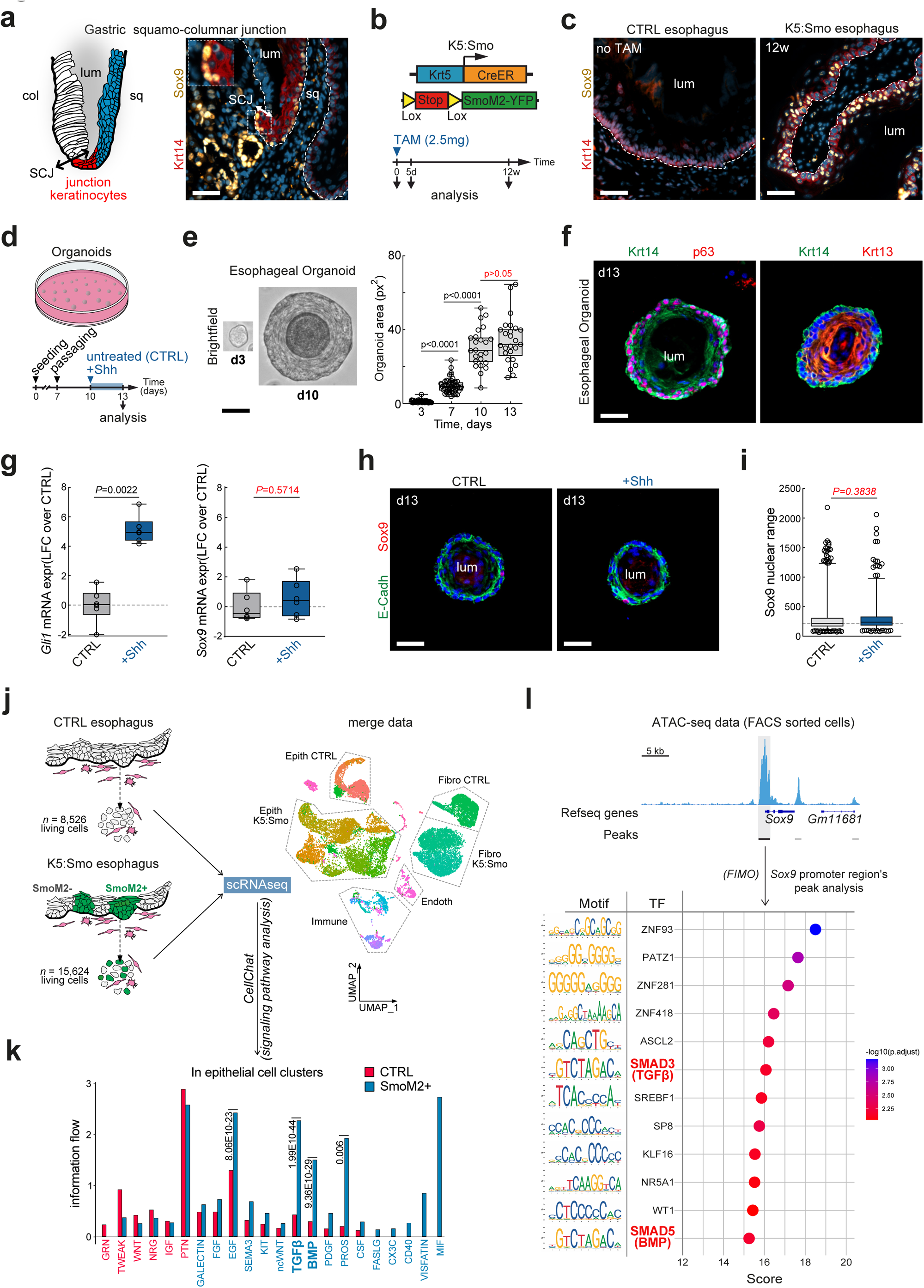
Lack of Direct Regulation of Sox9 Expression in Mouse Esophageal Progenitors by the Hedgehog Pathway *ex vivo*. (a) Schematic representation of the squamo-columnar junction (SCJ) and immunostaining for Krt14 and Sox9 in junction keratinocytes of control mice. (b) Transgenic mouse model used in this study: Krt5CreER;RosaSmoM2 (K5:Smo) and experimental design. (c) Immunostaining for Sox9 in the esophagus of K5:Smo mice 12 weeks after TAM administration and in control mice (no TAM). These data demonstrate the absence of Sox9 expression in the esophageal epithelium and its upregulation following activation of the HH signaling pathway. (d) Experimental design. (e) Micrographs of esophageal organoids between 3 and 10 days after seeding, and optical section area measured over time between 3 and 13 days after seeding. (f) Immunostaining for Krt14, p63, and Krt13 in esophageal organoids cultured for 13 days *ex vivo*. (g) Expression of *Gli1*, the HH target gene and *Sox9* mRNA measured by qPCR in esophageal organoids treated with Shh (*n* = 6). Data are represented as box plots (min to max) and expressed as the log2 fold change (LFC) relative to the control condition (untreated, *n* = 6). *P*-values were calculated using the Mann-Whitney test. (h) Immunostaining for E-Cadherin and Sox9 in esophageal organoids treated with Shh and in the control condition. (i) Boxplot summarizing Sox9 protein expression measured by immunofluorescence in esophageal organoids treated with Shh and in the control condition. *P*-value was calculated using t-test. (j) Experimental design of the single-cell RNA sequencing analysis from control and K5:Smo esophagi. Data were merged and visualized using UMAP. (k) Barplot representing enriched signaling pathways in epithelial cells of control mice (red) and in SmoM2+ epithelial cells of K5:Smo mice (blue). *P*-values were calculated using the Wilcoxon test with a significance threshold of 0.01. (l) Dotplot illustrating the ATAC-seq profile of *Sox9* promoter region in esophageal epithelial cells. Putative binding motifs for several transcription factors, including Smads, are found in the Sox9 transcription initiation site. Nuclear staining is represented in blue. Scale bars represent 50 µm. TAM, Tamoxifen administration; Col, columnar; Sq, squamous; SCJ, squamo-columnar junction; lum, lumen; CTRL, control; Shh, Sonic Hedgehog, K5:Smo; K5:Smo mice 12 weeks after TAM; dash lines represent the basal lamina.

To determine whether Sox9 expression is directly modulated by the HH pathway in esophageal mucosa, we utilized organoid cultures of mouse esophageal keratinocytes. When cultivated in a keratinocyte-defined medium, these organoids grow for 10 days and then reach a plateau (Fig. 1d, e). Until day 13 in culture, the expression patterns of the transcription factor p63 as well as the esophageal cell differentiation marker Krt13, are similar in the organoids *in vitro* and the esophagi *in vivo* (Fig. 1f and Supp. Fig. 1h, i). Thus, the organoids recapitulate the normal process of squamous differentiation and can be used to study the mechanisms that may modify this process.

In esophageal organoids, recombinant Sonic Hedgehog (Shh) protein, which effectively activates the HH pathway as shown by the significant upregulation of the HH target gene *Gli1* after a three-day treatment, fails to induce any significant variation in the level of *Sox9* (Fig. 1g). The same result was observed at the protein level by immunostaining (Fig. 1h, i). These results suggest that the upregulation of Sox9 triggered by the activation of the HH pathway *in vivo* is indirect and may require a relay from the microenvironment that is absent from the organoid culture condition.

To determine which cues from the microenvironment may regulate Sox9 expression, we used single-cell RNA sequencing (scRNA-seq) data of 15,624 living cells from K5:Smo mice 12 weeks after tamoxifen administration and merged them with data of 8,526 living cells from CTRL mice (Fig. 1j and Supp. Fig. 1j-l). Characterization of the signaling pathway network using *CellChat* ^7^ revealed that the Epidermal growth factor (EGF), Transforming growth factor beta (TGF-β), Bone morphogenetic protein (BMP), and protein S (PROS) pathways are more active in epithelial cells following HH pathway reactivation *in vivo* (Fig. 1k). To identify transcription factors and associated pathways that may directly regulate Sox9 expression in esophageal keratinocytes, we analyzed the open chromatin region in *Sox9* promoter using ATAC-sequencing data (Assay for Transposase-Accessible Chromatin sequencing) from FACS-sorted wild-type mouse esophageal epithelial cells^6^ (Fig. 1l). Using FIMO (Find Individual Motif Occurrences) ^8^, we identified several putative binding motifs in the *Sox9* promoter-associated peak. Notably, we identified Smad5 and Smad3 binding sequences upstream of *Sox9* (Fig. 1l and Table S1), suggesting that the TGF-β/BMP pathways may constitute a relay for the HH pathway that regulates *Sox9* transcription. Interestingly, the absence of Gli-binding motifs in this region further confirms that the HH pathway does not directly regulate Sox9 expression.

### Activation of the Hedgehog pathway in esophageal mucosa promotes modifications of the microenvironment that activate the TGF-β pathway *in vivo*

To determine whether HH pathway activation stimulates TGF-β signaling in esophageal cells, we investigated the level of Smad2 phosphorylation in the esophagus of control and K5:Smo animals (Fig. 2a, b). Whereas Phospho-Smad2 was found exclusively in suprabasal cells in control condition, it was upregulated in basal cells from K5:Smo esophagi, thus demonstrating that the HH pathway stimulates TGF-β pathway activation in esophageal progenitors *in vivo* (Fig. 2a, b and Supp. Fig. 2a). Interestingly, Phospho-Smad2 was detected already 5 days after TAM administration, at the time Sox9 appears in epithelial cells.

**Fig. 2.**
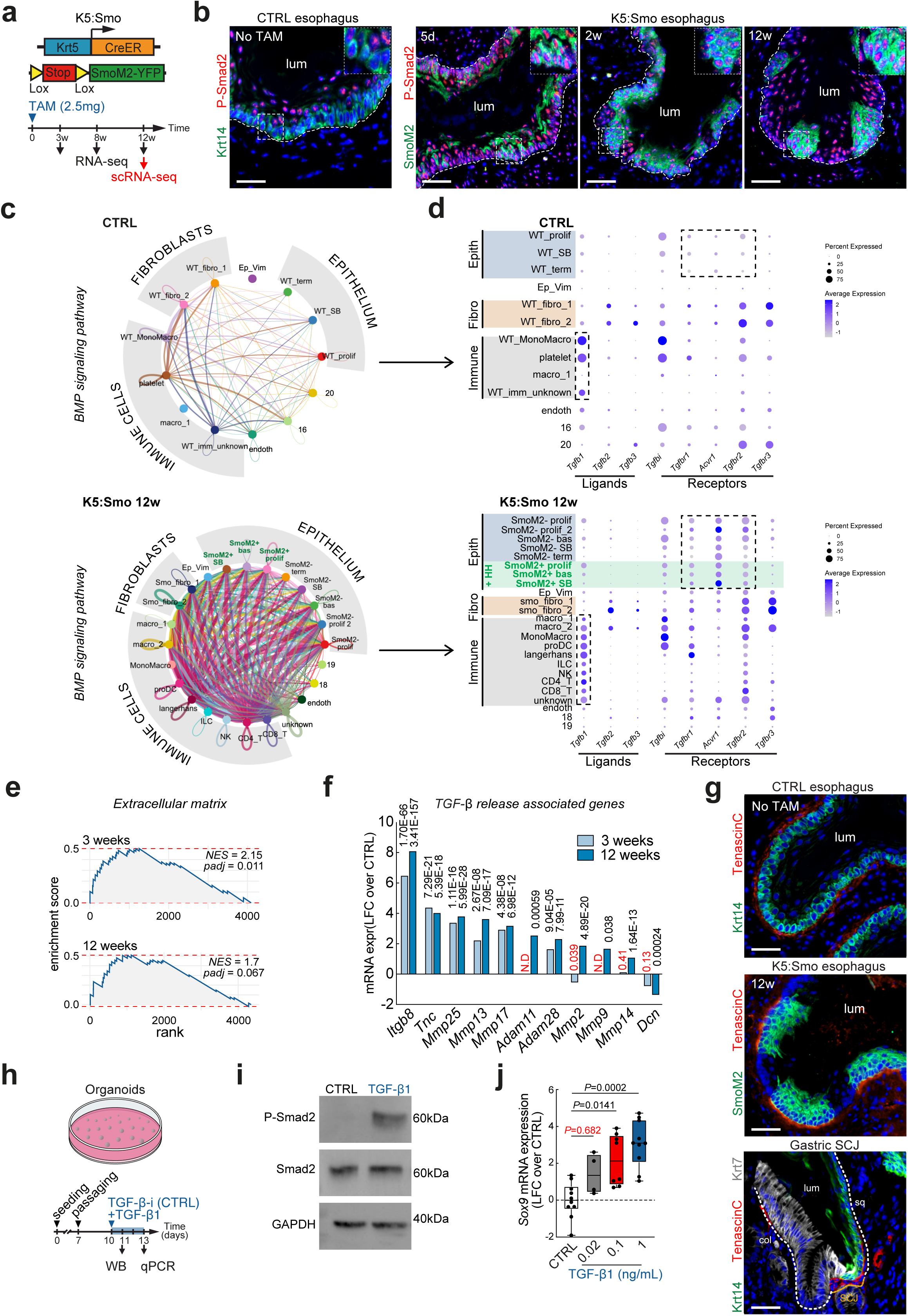
Hedgehog pathway stimulates TGF-β1/Smad2 pathway activation in esophageal cells *in vivo*. (a) Experimental design. (b) Immunostaining for P-Smad2 and Krt14 in control mice and for P-Smad2 and SmoM2-YFP in K5:Smo mice at 5 days, 2 weeks and 12 weeks after TAM. These data demonstrate that the TGF-β signaling pathway is enriched in esophageal epithelial cells following the activation of the HH pathway. (c) Circle plot showing the TGF-β mediated cell communication in esophageal cells from control and K5:Smo mice. (d) Dot plot illustrating the expression of transcripts related to the TGF-β pathway in esophageal cells from control and K5:Smo mice. (e) Gene set enrichment analysis (GSEA) of the significantly modified transcripts in SmoM2+ epithelial cells of K5:Smo mice 3 weeks (*n* = 4) and 12 weeks (*n* = 6) after TAM compared with control esophageal cells. (f) Expression of TGF-β release associated genes measured by RNA-seq in SmoM2+ epithelial cells of K5:Smo mice 3 weeks, 12 weeks after TAM compared to CTRL. These data show an upregulation of those transcripts in SmoM2+ epithelial cells compared to CTRL. (g) Immunostaining for Tenascin-C in the esophagus of control mouse in K5:Smo animal 12 weeks after TAM, as well as at the SCJ of control mouse. These data demonstrate that Tenascin-C is expressed by basal cells following HH pathway activation and at the SCJ of control mice. (h) Experimental design. (i) Expression of P-Smad2 and Smad2 measured by Western blot in esophageal organoids treated for 24 hours with TGF-β inhibitor (CTRL) or TGF-β. (j) Sox9 expression measured by qPCR in esophageal organoids treated with TGF-β inhibitor (*n* = 10), TGF-β 0.02 ng/ml (*n* = 4), TGF-β 0.1 ng/ml (*n* = 8) and TGF-β 1 ng/ml (*n* = 10). Data are represented as box plots (min to max) and expressed as *LFC* over CTRL. Nuclear staining is represented in blue. Scale bars represent 50 µm. CTRL, control; TAM, Tamoxifen administration; lum, lumen; Col, columnar; Sq, squamous; SCJ, squamo-columnar junction; dash lines represent the basal lamina.

To determine what stimulates TGF-β pathway in esophageal keratinocytes, we used scRNA-seq data of living cells from K5:Smo and from CTRL mice (Fig. 2c, d and Supp. Fig. 2b). Characterization of the TGF-β signaling pathway network using *CellChat* revealed that TGF-β signaling network is indeed upregulated in K5:Smo animals (Fig. 2c). These results also revealed that immune cells constitute the main source of TGF-β1 and showed an upregulation of TGF-β receptor 2 (*Tgfbr2*) and Activin receptor 1 (*Acvr1*) in basal progenitors from K5:Smo mice compared to control animals, probably increasing their sensitivity to TGF-β. (Fig. 2d).

The TGF-β pathway can be modulated in tissues by releasing latent TGF-β from the matrix ^9^. We therefore performed gene set enrichment analysis (GSEA) of RNA-seq data from K5:Smo esophageal progenitors compared to control keratinocytes. Interestingly, these data revealed that the gene set that is most significantly enriched in esophageal cells upon HH pathway activation is related to extracellular matrix (ECM) organization (Fig. 2e). We looked at the expression of ECM components in RNA-seq profiles of K5:Smo and control mice and found that Integrin β8 (*Itgb8*), Tenascin-C (*TnC*) and several metallopeptidases (*Mmp13*, *Mmp17*, *Mmp2*, and *Mmp9*) are significantly upregulated in esophageal epithelial cells in which the HH pathway is activated (Fig. 2f). An increase in Tenascin-C expression is compatible with a release of latent TGF-β and the subsequent activation of this pathway in the surrounding cells. Co-immunostaining for Tenascin-C and SmoM2 confirmed preferential expression of Tenascin-C in the ECM facing the SmoM2+ clones as well as in the ECM facing squamous SCJ progenitors (Fig. 2g and Supp. Fig. 2c, d). Together, these results suggest that the HH pathway activation in esophageal cells appears to promote the TGF-β pathway activation in the epithelium mostly by increasing the release of latent TGF-β trapped in the ECM and by increasing the expression of TGF-β receptors.

Since Matrigel contains a variable amount of TGF-β, we added a TGF-β receptor inhibitor (TGF-β-i) in our culture medium under control conditions to avoid basal Smad2 phosphorylation (Fig. 2h). As expected, treating cells with TGF-β1 induces the phosphorylation of Smad2 (Fig. 2i). In line with our hypothesis, we observed that increasing doses of TGF-β1 induced a progressive upregulation of *Sox9* in esophageal cells (Fig. 2j). In esophageal progenitors, Sox9 is therefore a target of the TGF-β/Smad2 pathway.

### The Bone morphogenetic protein signaling pathway potentiates the impact of TGF-β on Sox9 expression in esophageal progenitors *in vivo*

To determine whether the BMP pathway is activated in esophageal cells following HH pathway activation *in vivo*, we analyzed bulk RNA-seq data of FACS-sorted epithelial cells from wild-type and SmoM2+ animals 3, 8, and 12 weeks after TAM administration (Fig. 3a). GSEA of these data showed that transcripts associated with the BMP pathway tend to increase over time in cells in which the HH pathway is activated, but without reaching statistical significance (Fig. 3b). Nonetheless, analysis of these RNA-seq data revealed that some BMP pathway-related genes such as *Bmp1*, *Bmp6*, and *Bmp7* are significantly upregulated in keratinocytes 8 weeks after TAM administration and later (Fig. 3c).

**Fig. 3.**
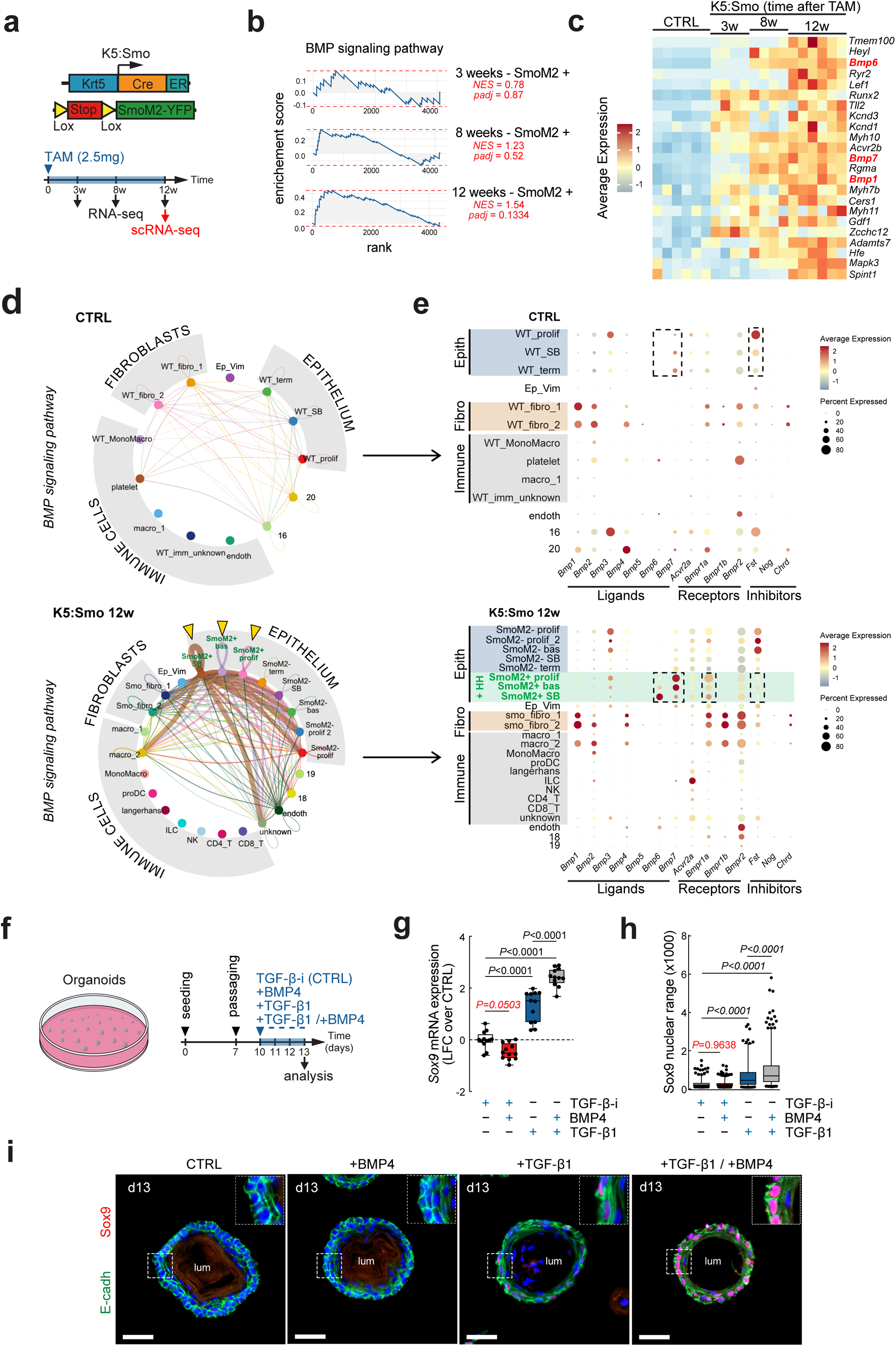
Hedgehog pathway activation in esophageal progenitors progressively induces a Bmp7 autocrine loop *in vivo*. (a) Experimental design. (b) Gene Set Enrichment Analysis (GSEA) of the significantly modified transcripts in SmoM2+ epithelial cells of K5:Smo mice 3 weeks (*n* = 4), 8 weeks (*n* = 4), and 12 weeks (*n* = 6) after TAM compared with control esophageal cells. (c) Heatmap representing the mRNA expression of the BMP signaling-related transcripts in control epithelial cells (*n = 6*) and in SmoM2+ epithelial cells of K5:Smo mice 3 weeks (*n* = 4), 8 weeks (*n* = 4), and 12 weeks (*n* = 6) after TAM. (d) Circle plot showing the BMP mediated cell communication in esophageal cells from control and K5:Smo mice. (e) Dot plot illustrating the expression of transcripts related to the BMP pathway in esophageal cells from control and K5:Smo mice. (f) Experimental design. (g) *Sox9* mRNA expression measured by qPCR in organoids treated with TGF-β inhibitor (*n* = 12), BMP4 (*n* = 12), TGF-β1 (*n* = 12) or a combination of TGF-β1 and BMP4. Data are represented as box plots (min to max) and expressed as LFC over TGF-β inhibitor condition. *P*-values were calculated using ANOVA followed by Tukey’s multiple comparison. (h) Boxplot (min to max) summarizing Sox9 protein expression measured by immunofluorescence in esophageal organoids treated for 4 days with TGF-β inhibitor, BMP4, TGF-β1 or a combination of TGF-β1 and BMP4. *P*-values were calculated using ANOVA followed by Tukey’s multiple comparison. (i) Immunostaining for E-Cadherin and Sox9 in esophageal organoids treated with TGF-β inhibitor, BMP4, TGF-β1 or a combination of TGF-β1 and BMP4. Nuclear staining is represented in blue. Scale bars represent 50 µm. lum, lumen. TAM, tamoxifen administration.

We then investigated the BMP signaling pathway using *CellChat* on scRNA-seq data (Fig. 3d, e). These data revealed that the BMP signaling network is modified in K5:Smo animals. Under control conditions, fibroblasts are the primary source of BMP ligands (*Bmp1*, *Bmp2*, and *Bmp4*) (Fig. 3e). In K5:Smo animals, we did not detect an upregulation of BMP ligands in fibroblasts but observed an increase in the level of *Bmp7* expression in the SmoM2+ clusters of epithelial cells (Fig. 3e and Supp. Fig. 3a-c). We also measured a significant upregulation of the Bmp receptor 1a *(Bmpr1a)* in esophageal keratinocyte clusters, suggesting that HH pathway activation may lead to an autocrine BMP loop in esophageal progenitors. In the same line, we observed a downregulation of the BMP repressor Follistatin (*Fst*) at the mRNA level in the SmoM2+ clusters of epithelial cells compared to control conditions (Fig. 3E and Supp. Fig. 3a-c). Bulk RNA-seq data showed that *Fst* downregulation and *Bmp7* upregulation occur 8 weeks after tamoxifen administration (Supp. Fig. 3a). These data suggest that in our model, the activation of the HH pathway in the epithelium may promote the progressive activation of an autocrine BMP loop.

To challenge the role of Smad1/5 in Sox9 transcription regulation, we treated esophageal organoids with BMP4. Using Western blot, we confirmed that BMP4 induces Smad1/5 phosphorylation, whereas TGF-β1 does not (Supp. Fig. 3d). Intriguingly, in these organoids, BMP4 did not promote Sox9 expression on its own, but further increased Sox9 expression in TGF-β1-treated cells (Fig. 3g). Immunostaining confirmed that BMP4 alone does not directly stimulate Sox9 expression, whereas TGF-β1 does. In addition, BMP4 potentiates the impact of TGF-β1 on Sox9 protein expression (Fig. 3h, i and Supp. Fig. 3e). Hence, our data suggest that the BMP pathway is not instrumental in inducing Sox9 expression in esophageal cells but would rather modulate its expression level.

### The Transforming growth factor-β and the Bone morphogenetic protein signaling pathways are complementary in the regulation of esophageal cell reprogramming *in vitro*

To assess whether TGF-β1, BMP4, or both modulate esophageal cell plasticity, we investigated the genome-wide consequences of a three-day treatment with each agonist alone or in combination (Fig. 4a). Consistent with our previous results on Sox9 expression, transcriptomic analysis revealed that the combination of TGF-β1 and BMP4 had the greatest impact on esophageal cells, although TGF-β1 alone had a strong impact as well (Fig. 4b). On the opposite, BMP4 alone appeared to have a more limited impact on esophageal cells RNA profile (Fig. 4b and Supp. Fig. 4a). We performed GSEA analysis to identify the biological functions modified upon TGF-β1/BMP4 co-treatment and found that expression of squamous differentiation-related genes was inhibited while "*extracellular matrix*" and "*digestive system development*" related genes were promoted (Fig. 4c). By looking at selected genes, we confirmed that the genes involved in squamous differentiation such as *Lce1b*, *Lce1j*, *Lce1d*, *Krt4*, *Krt15*, and *Krt13* were significantly downregulated while markers of columnar epithelium such as *Krt7*, *Krt8*, *Krt20*, and several mucins were upregulated (Fig. 4d). We looked at the transcripts modified by TGF-β1 or BMP4 alone and found that while BMP4 specifically represses expression of early squamous differentiation markers (*Krt4* and *Krt13*), TGF-β1 promotes expression of metaplasia markers such as *Krt7*, *Krt8*, *Krt20*, or *Sox9* (Supp. Fig. 4b). By immunofluorescence, we could appreciate that esophageal organoids treated with TGF-β1 and BMP4 for 3 days remained p63 positive but were characterized by Sox2 downregulation and Krt7 upregulation (Fig. 4e). This transcription factor expression pattern (p63-positive / Sox9-positive / Sox2-low) was reminiscent of SCJ keratinocytes (Fig. 4f). To determine to what extent TGF-β1 and BMP4 may promote a junction-like phenotype, we used bulk RNA-seq and compared the transcripts enriched in TGF-β1/BMP4-treated organoids to the transcripts enriched in junction keratinocytes. These analyses showed that about 20% of the genes enriched in junction keratinocytes are upregulated by TGF-β1 and BMP4 in just three days, while 20% of the transcripts enriched in normal keratinocytes are downregulated (Fig. 4g). In the same manner, about 20% of the genes deregulated by TGF-β1/BMP4 are common with transcripts deregulated in HH-stimulated esophageal progenitors *in vivo*, suggesting that these pathways could act as a relay for the HH pathway to promote cell reprogramming *in vivo*. (Supp. Fig. 4d). These data highlight the impact of the TGF-β/BMP signaling pathway on esophageal cell fate and point to a key role of epithelial-stromal crosstalk in cell plasticity. Nonetheless, additional mechanisms may participate in the regulation of Sox9 expression and esophageal cell plasticity.

**Fig. 4.**
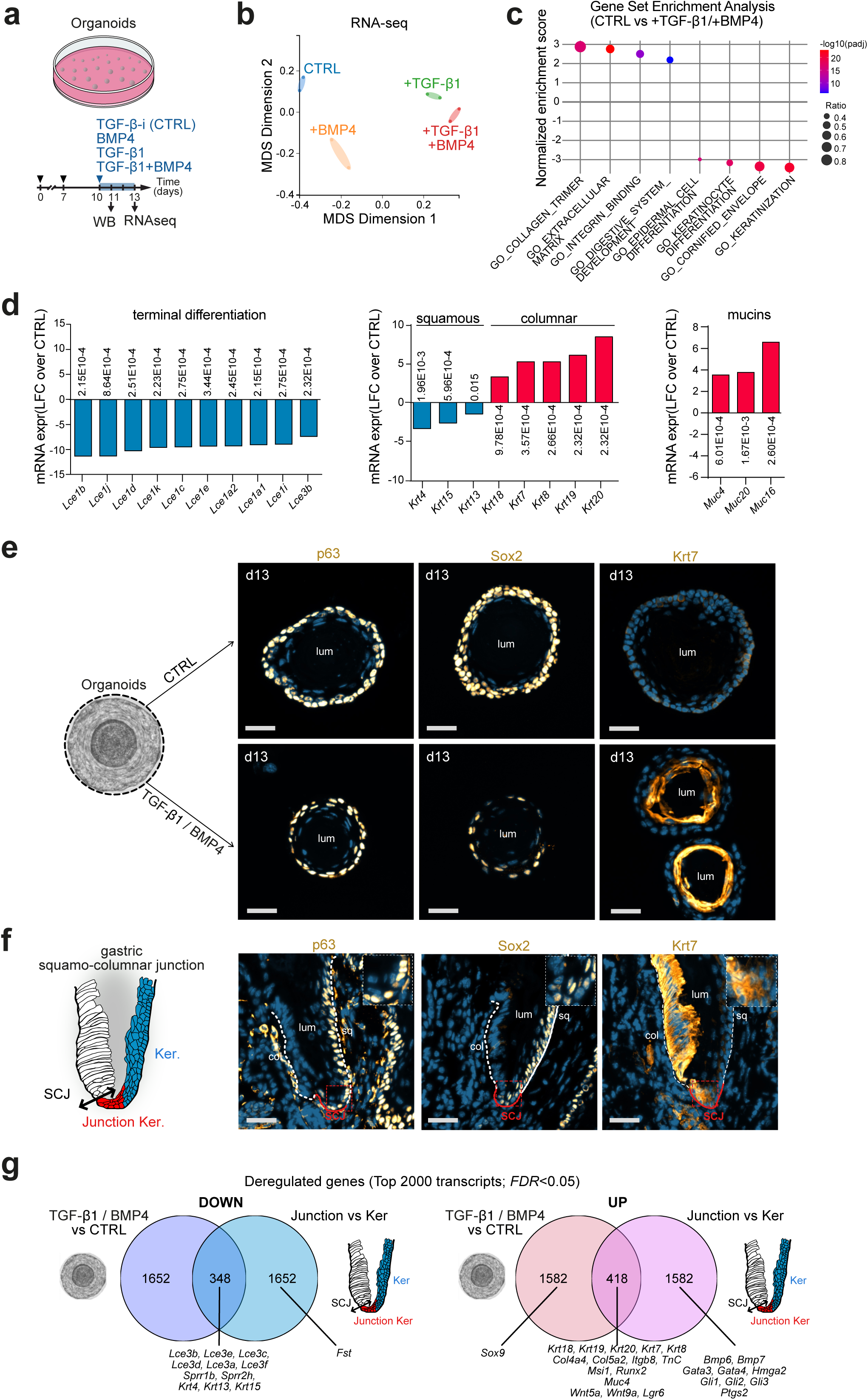
TGF-β/BMP signaling pathway activation in esophageal progenitors *ex vivo* partially recapitulate the phenotype of Hedgehog-stimulated cells *in vivo*. (a) Experimental design. (b) MDS plot representing mRNA sequencing data from CTRL organoids treated with TGF-β inhibitor (CTRL, *n = 2*), BMP4 (*n* = 2), TGF-β1 (*n* = 2) or a combination of TGF-β1 and BMP4 (*n* = 2). (c) Dotplot representing a selection gene ontology (GO) obtained after Gene Set Enrichment Analysis (GSEA) of significantly modified transcripts in esophageal organoids treated with TGF-β1 + BMP4 compared to CTRL. (d) mRNA expression of transcripts measured by RNA-seq in esophageal organoids treated with TGF-β1 + BMP4 compared to CTRL. The data show an upregulation of genes related to the columnar fate and a downregulation of genes associated with the terminal differentiation of squamous epithelial cells. (e) Immunostaining for p63, Sox2, or Krt7 in esophageal organoids treated with TGF-β1 + BMP4 compared to CTRL. (f) Immunostaining for p63, Sox2 and Krt7 at the SCJ from CTRL mice. (g) Venn diagrams comparing the top 2000 significantly deregulated transcripts in organoids treated with TGF-β1 + BMP4 compared to CTRL and in the squamous part of the SCJ (Krt7+ squamous cells) compared to normal esophageal epithelium. These data show that 17.4% of downregulated genes and 20.9% of upregulated genes are common between the TGF-β1+BMP4 -treated organoids and the Krt7+ squamous cells from the SCJ. Nuclear staining is represented in blue. Scale bars represent 50 µm. CTRL, control; Col, columnar; sq, squamous; SCJ, squamo-columnar junction; lum, lumen; dash lines represent the basal lamina.

### Ibuprofen prevents Hedgehog-driven Sox9 expression in vivo

Our bulk RNA-seq and scRNA-seq data revealed that *Ptgs2* coding for the cyclooxygenase-2 (Cox-2), is one of the most upregulated genes in esophageal cells following HH pathway activation and its expression increases with esophageal cell reprogramming (Fig. 5a-c). Previous study showed that COX-2 expression is significantly correlated with exposure of the distal esophagus to acid reflux ^10^. In addition, non-steroidal anti-inflammatory drugs (NSAIDs) may delay the development of metaplasia and adenocarcinoma in the esophagus ^11,12^. We therefore hypothesized that NSAIDs such as Ibuprofen, which inhibit Cox-2, may alter esophageal cell plasticity. To address this question, we compared esophagi from K5:Smo mice treated with ibuprofen to untreated animals (Fig. 5d). Strikingly, Sox9 protein is virtually absent from ibuprofen-treated K5:Smo mice, whereas it is expressed in SmoM2+ epithelial cells from untreated K5:Smo animals (Fig. 5e and Supp. Fig. 5a). The Sox9 target gene Krt7 was also absent from ibuprofen-treated SmoM2+ cells (Fig. 5e and Supp. Fig. 5a). These data therefore clearly show that Ibuprofen may alter Sox9-driven esophageal cell reprogramming *in vivo*.

**Fig. 5.**
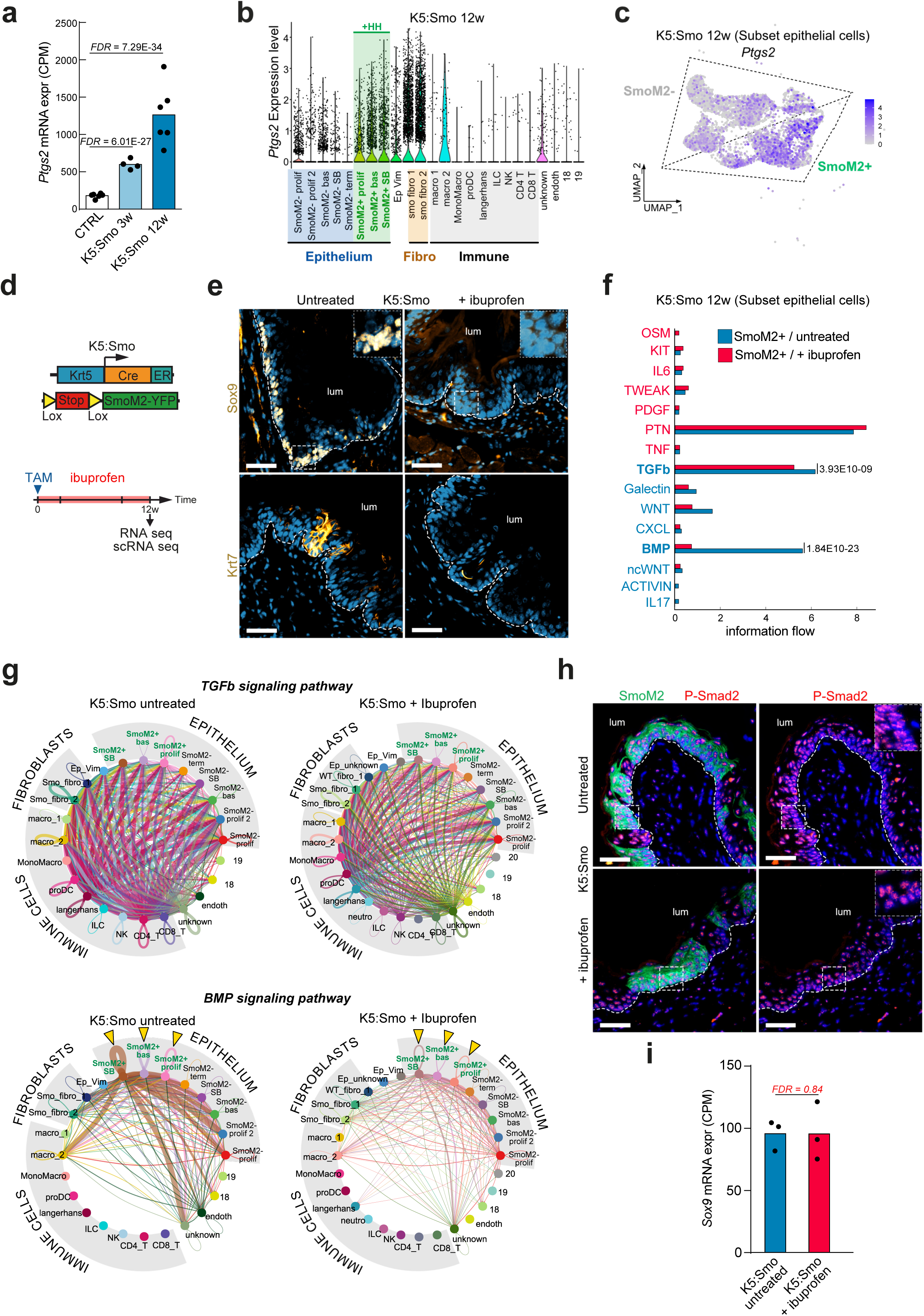
Ibuprofen inhibits Hedgehog-induced Sox9 expression in esophageal progenitors *in vivo*. (a) *Ptgs2* mRNA expression measured by RNA-seq in control esophageal epithelial cells (*n =* 6) and in SmoM2+ esophageal epithelial cells of K5:Smo mice 3 (*n = 4*) and 12 weeks (*n = 6*) post-TAM. (b) Violin plots showing *Ptgs2* mRNA expression in scRNA seq data of K5:Smo mice. *Ptgs2* is virtually absent in SmoM2-negative epithelial cells and upregulated in SmoM2-positive epithelial cells. (c) Feature plot representing *Ptgs2* expression in the subset of epithelial cell clusters of K5:Smo mice. (d) Experimental design. (e) Immunostaining for Sox9 and Krt7 in untreated and ibuprofen-treated 0.6g/l K5:Smo mice 12 weeks post-TAM. These pictures show that Sox9 and Krt7 are downregulated after ibuprofen treatment. (f) Barplot representing enriched signaling pathways in SmoM2-positive esophageal epithelial cells from untreated (blue) or ibuprofen-treated (red) K5:Smo mice. *P*-values were calculated using Wilcoxon test with a significance threshold of 0.01. These data predicts that BMP signaling is strongly downregulated in SmoM2-positive epithelial cells after ibuprofen treatment. (g) Circle plot showing the TGF-β and BMP mediated cell communication in esophageal cells from untreated or ibuprofen-treated K5:Smo mice 12 weeks after TAM. (h) Immunostaining for P-Smad2 and SmoM2-YFP in untreated and ibuprofen-treated K5:Smo mice. These pictures show that Smad2 phosphorylation in K5:Smo epithelial cells is not affected by ibuprofen treatment. (i) *Sox9* mRNA expression in SmoM2+ esophageal epithelial cells in untreated and in ibuprofen-treated K5:Smo mice. Those data show that *Sox9* mRNA expression is not affected by ibuprofen treatment. Nuclear staining is represented in blue. Scale bars represent 50 µm. CTRL, control; lum, lumen; TAM, Tamoxifen administration; dash lines represent the basal lamina.

To determine whether Ibuprofen blocks Sox9 appearance in esophageal cells by modulating epithelial-stromal interactions, we studied signaling pathway networks in epithelial cells using scRNA-seq. These data showed that Ibuprofen treatment leads to a slight decrease in TGF-β signaling by immune cells and inhibits the BMP autocrine loop in epithelial cells (Fig. 5f, g). In the same line, we measured Smad2 phosphorylation levels by immunofluorescence and did not detect any notable impact of ibuprofen treatment (Fig. 5h and Supp. Fig. 5d). Importantly, we observed that while Sox9 protein is virtually absent from ibuprofen-treated K5:Smo esophagi (Fig. 5e), its RNA expression appeared unaltered, suggesting a cell-autonomous mechanism and a potential regulation at the protein-level (Fig. 5i).

### Ibuprofen directly targets esophageal progenitors to inhibit Sox9 protein expression

To determine whether Ibuprofen directly affects Sox9 expression in keratinocytes, we treated K5:Smo esophageal organoids with TGF-β1 and BMP4, with or without ibuprofen (Fig. 6a). Our protocol allows to cultivate both SmoM2-negative and -positive organoids from the same esophagus in the same well to decrease technical biases. Under control conditions, Sox9 protein is virtually absent from both SmoM2-negative and -positive organoids, demonstrating that the HH pathway, on its own, is not sufficient to induce Sox9 expression (Supp Fig.6a and Fig. 6c). In addition, TGF-β1/BMP4 induced Sox9 expression in both organoids, although its level was higher in SmoM2-positive organoids. Interestingly, the TGF-β1/BMP4-induced Sox9 protein was decreased in K5:Smo organoids treated with Ibuprofen, in both SmoM2+ and SmoM2-organoids (Fig. 6b, c). Ibuprofen also prevented expression of the Sox9 target gene Krt7 following TGF-β1/BMP4 treatment (Fig. 6d, e and Supp Fig. 6b). Whereas TGF-β1/BMP4-induced *Krt7* transcription was inhibited when organoids were treated with Ibuprofen, no modifications were observed in *Sox9* transcript levels, suggesting that Ibuprofen primarily affects Sox9 protein stability (Fig. 6e). In line with this observation, Ibuprofen decreases Sox9 protein expression levels without inhibiting Smad2 phosphorylation but with partial inhibition of Smad1/5 phosphorylation (Fig. 6f, g and Supp Fig.6 c-e). Finally, RNA-seq profiling of treated organoids revealed that ibuprofen prevents about 40% of the transcriptional changes induced by TGF-β1/BMP4 treatment (Fig. 6h). Together, these data suggest that Ibuprofen may directly inhibit Sox9 protein expression in esophageal progenitors, thereby stalling TGF-β/BMP-induced cell reprogramming.

**Fig. 6.**
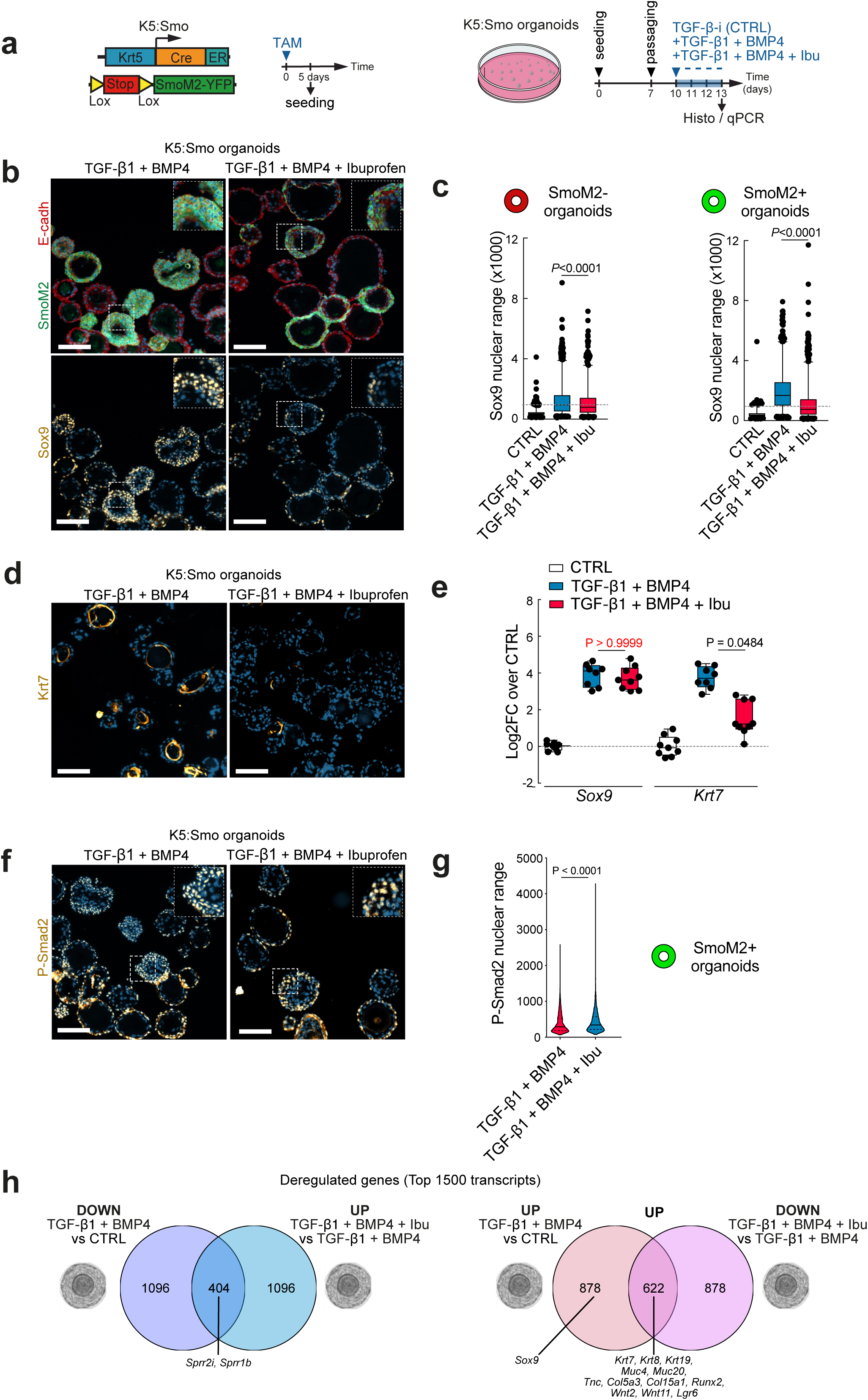
Ibuprofen inhibits TGF-β/BMP-stimulated Sox9 protein expression but not its mRNA expression in mouse esophageal progenitors *ex vivo*. (a) Experimental design. (b) Immunostaining for SmoM2-YFP, E-cadherin and Sox9 in K5:Smo esophageal organoids treated with TGF-β1 + BMP4 or with TGF-β1 + BMP4 + ibuprofen. These pictures show Sox9 downregulation after ibuprofen treatment. (c) Boxplot (min to max) summarizing Sox9 protein expression measured by immunofluorescence in K5:Smo esophageal organoids (SmoM2 negative or SmoM2 positive organoids) treated with TGF-β inhibitor (CTRL), TGF-β1 +BMP4 or with TGF-β1 + BMP4 + ibuprofen. (d) Immunostaining for Krt7 in K5:Smo esophageal organoids treated with TGF-β1 + BMP4 or with TGF-β1 + BMP4 + ibuprofen. Pictures show a downregulation of Krt7 after ibuprofen treatment. (e) *Sox9* and *Krt7* mRNA expression in K5:Smo organoids treated with TGF-β inhibitor (CTRL) (*n* = 9), TGF-β1 +BMP4 (*n* = 8) or TGF-β1 + BMP4 + ibuprofen (*n* = 9), measured by qPCR. Data are represented as box plots (min to max) and expressed as LFC over CTRL. (f) Immunostaining for SmoM2 (YFP), P-Smad2 and P-Smad1/5 in K5:Smo organoids treated with TGF-β1 + BMP4 or with TGF-β1 + BMP4 + ibuprofen (g) Boxplot (min to max) summarizing Smad2 protein phosphorylation measured by immunofluorescence in K5:Smo esophageal organoid treated for 3 days with TGF-β1 + BMP4 or TGF-β1 + BMP4 + ibuprofen. (h) Venn diagrams comparing the top 1500 deregulated transcripts in wild-type organoids treated with TGF-β1 + BMP4 compared to TGF-β inhibitor (CTRL) and in TGF-β1 + BMP4 + Ibu compared to TGF-β1 + BMP4. These data show that about 40% of the genes upregulated by TGF-β1 + BMP4 treatment are downregulated by ibuprofen. In the same manner, 27% of the genes downregulated by TGF-β1 + BMP4 treatment are upregulated by ibuprofen. Nuclear staining is represented in blue. Scale bars represent 100 µm. CTRL, control; TAM, Tamoxifen administration.

### Ibuprofen inhibits Transforming Growth Factor beta-induced SOX9 expression in human esophageal cells ex vivo

To validate the data obtained in mouse esophageal cells, we tested the impact of TGF-β1/BMP4 on SOX9 expression in human esophageal epithelial cells, as well as the inhibitory effect of Ibuprofen. We grew human esophageal organoids by cultivating enzymatically digested esophageal biopsies in Matrigel (Fig. 7a and Supp Fig.7a, b). These organoids are composed of α6-INTEGRIN positive basal epithelial cells and several layers of KRT13 positive differentiated cells, thereby recapitulating the normal organization of the esophageal epithelium (Fig. 7b and Supp Fig. 7c). Similar to the data we observed in mouse, a three-day treatment with TGF-β1/BMP4 did not block P63 but induced SOX9 expression while decreasing SOX2 expression (Fig. 7c and Supp Fig. 7d-f). As observed in K5:Smo mouse esophageal organoids, Ibuprofen blocked TGF-β1/BMP4-induced SOX9 protein expression (Fig. 7d, e and Supp Fig. 7g). On the opposite, Ibuprofen did not block TGF-β1/BMP4-induced *SOX9* transcription (Fig. 7f). Importantly, the Cox-2 inhibitor Celecoxib had the same effect as Ibuprofen on TGF-β1/BMP4-induced SOX9 expression at both mRNA and protein levels (Fig. 7d-f). Together, the results obtained from both mouse and human models suggest that pharmacologically targeting Sox9 expression could be used to modify esophageal progenitor cell reprogramming.

**Fig. 7.**
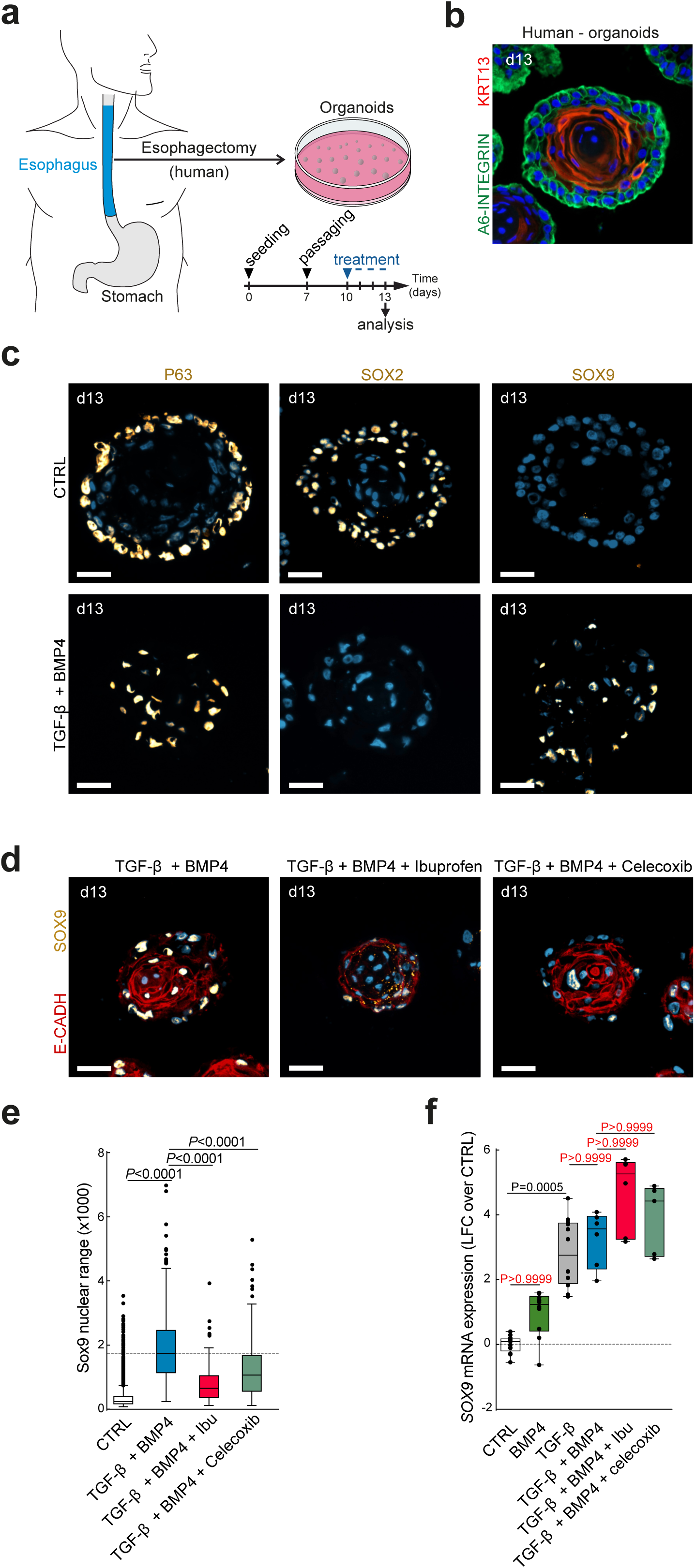
Ibuprofen inhibits TGF-β/BMP-stimulated SOX9 protein expression but not its mRNA expression in human esophageal progenitors. (a) Experimental design. (b) Immunostaining for A6-INTEGRIN and KRT13 in human esophageal organoids. (c) Immunostaining for P63, SOX2 and SOX9 in human esophageal organoids treated with TGF-β inhibitor (CTRL) or TGF-β1 +BMP4. (d) Immunostaining for E-CADH and SOX9 in human esophageal organoids treated with TGF-β1 + BMP4, TGF-β1 + BMP4 + ibuprofen or TGF-β1 + BMP4 + celecoxib. (e) Boxplot (min to max) summarizing Sox9 protein expression measured by immunofluorescence in human esophageal organoids treated with TGF-β inhibitor (CTRL), TGF-β1 + BMP4, TGF-β1 + BMP4 + ibuprofen or TGF-β1 + BMP4 + celecoxib. These data shows that COX-2 inhibitors induce SOX9 downregulation in human esophageal cells. (f) *Sox9* mRNA expression in human esophageal organoids treated with TGF-β inhibitor (*n* = 5), BMP4 (*n* = 4), TGF-β1 (*n* = 4), TGF-β1 + BMP4 (*n* = 2), TGF-β1 +BMP4 + ibuprofen (*n* = 3) or TGF-β1 +BMP4 + celecoxib (*n* = 2), measured by qPCR. Data are represented as box plots (min to max) and expressed as *LFC* over CTRL. Nuclear staining is represented in blue. Scale bars represent 50 µm. CTRL, control.

## Discussion

### Role of esophageal cell plasticity

There is evidence to suggest that the esophageal epithelium exhibits some degree of cellular plasticity in response to certain signals or stimuli. One example of this plasticity is the columnar metaplasia of the esophagus, where cells in the esophageal epithelium undergo a transformation from squamous to columnar-like cells ^6^. This process has been observed in response to chronic injury or inflammation, such as that caused by GERD or BE ^13^.

Lineage tracing has unambiguously demonstrated that a subset of keratinocytes from the SCJ can give rise to BE-like metaplasia *in vivo* ^3^. In the same line, the presence of the same mitochondrial mutations in squamous mucosa and BE suggests a common cellular origin for keratinocytes and metaplastic cells ^14^. Metaplasia can develop not only in the distal but also in the proximal esophagus, in the context of esophago-cardia resection ^13,15^. Lineage tracing experiments showed that IL-1β-induced BE-like metaplasia originates from Lgr5 progenitors ^16^, while surgically induced metaplasia in the mid-esophagus does not ^13^. Together, these results suggest that different cells, including keratinocytes, may contribute to metaplasia ^1^.

The SCJ is composed of undifferentiated cells, which have been described as an embryonic-like tissue that may participate in columnar metaplasia following chronic squamous epithelium injury ^17^. A recent study reported that adjacent squamous and columnar epithelial cells may both contribute to the formation of the SCJ cells in the human esophagus ^18^. Single-cell characterization of the mouse gastroesophageal junction development failed to identify a transcriptomic trajectory linking squamous progenitors to the transition epithelium ^19^. Hence, it remains unclear whether the transcommitment of squamous progenitors into the transition epithelium occurs under physiological conditions or whether it would be restricted to chronic reflux context ^13^. Lineage tracing should be used in both conditions to address this question. Our group used this method to show that the HH pathway reactivation in esophageal progenitors, triggered by GERD, can induce the transition of Krt5+ squamous progenitors into embryonic-like undifferentiated columnar cells ^6^. This work provided a molecular basis for esophageal cell plasticity and pointed to Sox9 as a crucial transcription factor involved in the process of squamous-to-columnar conversion *in vivo*. Several studies have identified transcription factors such as CDX1, CDX2, GATA6, HOX, KLF4, and SOX9 as being involved in the development of columnar metaplasia of the esophagus ^3,5,6,20–23^. Understanding how the microenvironment modulates the expression of such transcription factors is therefore important.

### Role of the microenvironment in esophageal cell plasticity

Modifications of the mesenchyme and subsequent evolution of morphogens in the microenvironment such as TGF-β, BMP, Wnt, and FGF (Fibroblast growth factor) are involved in the development of gastric epithelium and squamous epithelium from embryonic bipotent progenitors ^19^. In this line, our data point to the crucial role of the microenvironment in esophageal cell plasticity regulation by modulating Sox9 expression and its target genes. In epidermal cells, HH pathway activation through SmoM2 expression or *Ptch1* deletion takes several weeks before inducing Sox9 expression, pointing to an indirect regulation in this model as well ^24^. These data suggest that HH pathway activation does not induce only an intrinsic plasticity but also an extrinsic plasticity, i.e, induced by changes in the microenvironment ^25,26^. Such an extrinsic cell plasticity was already reported in epidermal cells ^27^ and in airway epithelial cells ^28^, but it was not described in esophageal epithelium *in vivo*. This might be important since extensive stromal remodeling may contribute to metaplasia development, as reported in the transition zone between endocervix and ectocervix ^29^.

Beyond these general principles about extrinsic regulation, recent evidence suggests that physiological cues may create a regionalization in esophageal cell plasticity. First, keratinocytes from the SCJ demonstrate enhanced plasticity compared to those from other regions^3,13^. Within the esophagus itself, a gradient of Troy expression exists along the cephalo-caudal axis, indicating that the Wnt pathway may be differentially stimulated in the distal esophagus compared to the proximal segment ^30^. In addition, our group has recently shown that esophageal squamous progenitors can give rise to appendages called taste buds specifically in the cervical segment of the esophagus ^31^. Taste buds are non-squamous histological appendages involved in taste molecules detection. Their formation is tightly regulated by epithelial-mesenchymal crosstalk, which involves multiple signaling pathways, including BMP, Wnt, and Hedgehog ^32–34^. Together, these data suggest that modifications in the epithelial-mesenchymal crosstalk could modify esophageal progenitors plasticity and thus be a critical component of tissue repair and/or metaplasia development.

Identifying specific cues associated with these physiological and pathological cellular processes could be important. Modulating cell plasticity may represent an interesting therapeutic approach by repressing Sox9-driven transcommitment of esophageal progenitors and promoting their normal squamous differentiation, which could improve current treatments such as ablative strategies in precancerous metaplasia.

### Role of the TGF-β pathway in esophageal cell plasticity

We identified putative binding motifs for Smad3 and Smad5 in the open chromatin region upstream of the *Sox9* transcription initiation site, pointing to the potential role of the TGF-β/BMP signaling pathway in Sox9 regulation.

Our data show that TGF-β1 can stimulate Sox9 expression in esophageal progenitor cells *ex vivo*. Some studies have concluded that the canonical TGF-β receptors TGFΒR1 and TGFΒR2 are sufficient for phosphorylation of both SMAD2/3 and SMAD1/5 ^35,36^. Our results show that TGF-β1 does not induce Smad1/5 phosphorylation in esophageal cells. In addition, BMP4-mediated Smad1/5 phosphorylation is not sufficient to promote Sox9 expression in esophageal progenitors *in vitro*.

In several epithelia, including the esophageal mucosa, the TGF-β pathway regulates cell differentiation and its inhibition enhances the renewal of basal progenitors ^37^. In addition, TGF-β inhibition is critical for specification of p63-positive esophageal progenitors cells from p63-negative anterior foregut, as suggested by the study of human pluripotent stem cells ^38^. Recent studies showed that mesenchymal cells trigger a TGF-β-dependent fetal-like reprogramming in damaged intestinal epithelium ^39,40^. Interestingly, in esophageal organoids, TGF-β promotes a transcriptomic program resembling the one found in SCJ-progenitors and in HH-stimulated cells, which have both been reported to share similarities with embryonic esophagus ^6,17^. *In vivo*, we observed that upon HH pathway reactivation, the TGF-β signaling is globally overactivated in esophageal mucosa. This stimulation is associated with an increase in Sox9 protein expression specifically in the cells where the HH pathway is activated. *In vivo*, Smad2 phosphorylation does not promote Sox9 expression in SmoM2-negative cells, but it does *in vitro*. These results raise the question of the mechanisms inducing this dichotomy. One possibility is that the transcriptomic and epigenetic reprogramming induced by the HH pathway ^6^ modifies the Smad2 target genes and that growing esophageal cells in culture may induce similar modifications. Addressing this question would require further experiments.

### Role of the BMP pathway in esophageal cell differentiation

A recent study highlighted the enrichment markers associated with HH and TGF-β/BMP signaling pathways in human BE biopsies ^41^. Moreover, BMP4 overexpression in foregut keratinocytes leads to metaplasia at the SCJ, and together with CDX2, plays a role in intestinalization of metaplasia ^13^. The conditional deletion of Noggin, a small secreted protein that acts antagonizes BMP signaling, results in the development of a BE-like neo-columnar epithelium at the SCJ as well ^41^. The inhibition of BMP2/BMP4 blocked the neo-columnar lineage while promoting the development of a squamous epithelium. This study therefore suggests that both BMP2 and BMP4 may be attractive therapeutic targets for the treatment of BE ^13,42^. In the same line, BMP pathway inhibition could also prevent the transition from metaplasia to adenocarcinoma in an animal model ^43^.

It has already been reported that in a tissue reconstruction model, the HH pathway may induce Sox9 expression in a Bmp4 dependent manner in esophageal cells ^5^. However, we failed to recapitulate this effect of Bmp4 on Sox9 expression in mouse esophageal cells *ex vivo*. Our results did not show any increase in the BMP ligand production by fibroblasts following the reactivation of the Hedgehog-pathway, but rather the emergence of an autocrine Bmp7 loop in esophageal progenitors combined with the downregulation of Fst, which may reinforce the Bmp7 autocrine loop as observed in lingual taste buds ^32^. Fst is also a well-known inhibitor of Activin, a ligand of the TGF-β / BMP pathway that may play a role in metaplasia development in the esophagus ^44^. Modifications of the BMP signaling pathway in esophageal keratinocytes may therefore participate in the late phase of reprogramming.^5,6,20,32,44^

We observed that HH-induced Sox9 upregulation in esophageal cells *in vivo* occurs before we detect evidence of BMP pathway activation, suggesting that BMP is not the main driver of *Sox9* transcription. Still, our results show that BMP may play a role in the inhibition of the squamous differentiation program that is important for the step of squamous-to-columnar conversion observed during transcommitment^6^. The highest degree of cell reprogramming is indeed observed when cells were treated with a combination of TGF-β1 and BMP4.

Together, our data highlight the crucial role of the microenvironment in regulating esophageal cell plasticity, driven by a sum of cues coming from different cell types and several signaling pathways. Characterizing the remodeling of epithelial-mesenchymal interactions during the early stages of metaplasia development and its progression to dysplasia will be critical for a better understanding of this condition.

### Pharmacological regulation of esophageal cell plasticity

Non-steroid anti-inflammatory drugs (NSAIDs) are cyclooxygenase inhibitors, utilized as anti-inflammatory, analgesic and antipyretic medication. It has been reported that NSAIDs may inhibit, or at least delay, the development of BE and its progression toward cancer in animal models ^11,12^ and humans ^45^. We therefore wondered whether NSAIDs, such as Ibuprofen, could alter esophageal cells plasticity and ultimately keratinocytes transdifferentiation. In line with this hypothesis, we observed that chronic treatment with Ibuprofen was associated with a complete blockage of Sox9 protein expression in esophagus epithelium following HH pathway activation. In addition, Ibuprofen inhibits TGF-β1 and BMP4 induced Sox9 expression in esophageal K5Smo organoids. This suggest that its impact *in vivo* may be the consequence of a modification in the epithelium-mesenchymal interactions combined with a direct effect on keratinocytes.

Ibuprofen is not a selective drug and may impact several pathways including NRF2 ^46^, which has been reported to directly regulate Sox9 expression in chondrocytes ^47^. Our data on human organoids confirmed the impact of Ibuprofen on SOX9 protein expression. In addition, the use of Celecoxib, a Cox-2 specific inhibitor, recapitulated the effect of Ibuprofen on TGF-β1/BMP4 induced SOX9 expression in human esophageal cells. These data therefore point to the central role of Cox-2 in the regulation of Sox9 protein expression. Cox-2 has been reported to regulate Sox9 expression *in vitro* ^48^. Several mechanisms have been associated to Sox9 protein stabilization ^49–52^, yet these mechanisms are poorly understood. Further experiments will be required to determine how HH and TGF-β/BMP pathways regulate Sox9 protein expression in esophageal progenitors.

In conclusion, our study highlights the central role of the TGF-β/BMP signaling pathways in Sox9 expression regulation and subsequent esophageal progenitor cell plasticity in mouse and human. Therapeutic strategy targeting esophageal cell plasticity to maintain progenitors in their squamous lineage might be an option as a complementary approach for the treatment of BE in combination with ablative therapies such as submucosal endoscopic dissection or cryoablation. The results we obtained with Ibuprofen treatment on Sox9 expression is a proof of concept that cell plasticity could be challenged *in vivo* by pharmacological means.

## Acknowledgment

We thank the FACS core facility and the animal house facility of Université libre de Bruxelles (ULB) for their help. Sequencing was performed at the Brussels Interuniversity Genomics High Throughput core (www.brightcore.be) by Dr Frédérick Libert and Anne Lefort. B.B. is an investigator of Fonds de la Recherche Scientifique (FNRS) at ULB. L.D. is supported by a fellowship from the TELEVIE (n°40007654). B.D. is supported by *Fonds pour la formation à la recherche dans l’industrie et dans l’agriculture* (FRIA). This work was supported by Fondation contre le Cancer (FCC/ULB 2018-067 and 2022-164) and the FNRS (CDR n°400008574).

## Contribution

L.D, B.D and B.B designed, validated and conducted experiments. B.D and L.D performed single-cell RNA sequencing data analysis supervised by A.V.D and B.B; A.V.D provided advice on data analysis; F.C, L.C and L.V provided esophagus biopsies for the human organoids; M.L and M.I.G provided expertise in the organoid experiments. L.D, B.D and B.B conceived the project, supervised experiments, and wrote the manuscript. Funding acquisition and supervision of the project by B.B. Preparation of the manuscript by all authors.

## Methods

### Experimental model and subject details

The Krt5-CreERT2 knock-in mice (The Jackson Laboratory, Stock#029155, RRID:IMSR_JAX:029155) were crossed with the R26SmoM2 mice (The Jackson Laboratory, Stock#005130, RRID:IMSR_JAX:005130) in order to generate K5-CreERT2:R26SmoM2. 2.5 mg of tamoxifen resuspended in sunflower oil was injected intraperitoneally to K5-CreERT2:R26SmoM2 mice. Littermates of the same sex were randomly assigned to experimental groups. Mouse colonies were maintained under pathogen-free conditions in a certified animal facility in accordance with the European guidelines. All the experiments were approved by the ethical committee from the university and conform with regulatory standards (LA1230406 – projects 844N and 793N).

Human esophageal biopsies were obtained through the collaboration with the Anatomical Pathology Centre of the H.U.B. Normal esophageal mucosa was dissected in the upper margin of esophagectomy. This procedure was approved by the institutional Ethical Committee from the IJB, (protocol CE 3344 – MTA Beck_1077_IJB Sept 2021).

### Method details

For all experiments presented in this study, sample size was large enough to measure the effect size. No randomization and no blinding were performed in this study.

### Tissue digestion

Murine esophagi were dissected, minced and digested in 2 mg/mL of collagenase I (A&E scientific) during 1h, EDTA 5 mM during 20 min and trypsin 0.125% (A&E scientific) during 10 min. Human esophageal biopsies were minced and digested in 2.5 mg/mL of collagenase I (A&E scientific) during 30 min, EDTA 5 mM during 20 min and trypsin 0.25%, (A&E scientific) during 10 min. For both, incubations have been done on a rocking plate at 37°C.

For all the tissues, cells were then rinsed in PBS supplemented with 2% FBS and filtered through a 70 µm cell strainers.

### Murine and human spheroid culture

Murine esophageal keratinocytes from 2 different mouse models were used: wild-type (WT) or K5-CreER T2 :R26SmoM2. For the K5-CreER T2 :R26SmoM2 model, 2.5 mg TAM was injected to mice and these animals were sacrificed 5 days later. Murine or human esophagi were digested as described above. Protocol for spheroid culture was adapted from Kasagi et al., 2018. Murine single cell suspension was mixed with Matrigel (Corning®) and plated. Human single cell suspension was mixed with Growth Factor Reduced Matrigel (Corning®) and plated. After polymerization of the Matrigel by incubating at 37 °C for 10 min, medium (KSFM, Thermofisher, table S5) complemented with Rock inhibitor (Y-27632, 10 µM) was added. Medium was changed after 24 hours and then every 48 hours. Plates were maintained at 37°C in a 5% CO_2_ humidified incubator.

Organoids were splitted between day 5 and day 7 depending on their growth. Matrigel was disrupted with cold and sterile PBS. Organoids were centrifugated and resuspended in Matrigel and plated again. Organoids have been treated for72h for mRNA and histological analysis. Used agonists were the following: Shh (Biotechne, 100 ng/ml), recombinant mouse TGF-β1 (Biolegend, 1ng/ml), recombinant human TGF-β1 (Biolegend, 1ng/ml), recombinant mouse BMP-4 (Biotechne, 20 ng/ml), A83-01 (TGF-β inhibitor, Sigma-Aldrich®, 500 nM), ibuprofen (Sigma-Aldrich®, 500 µM).

### FACS isolation

Immunostaining was performed on single cell suspension using PE-conjugated anti-CD45 (1:500, BioLegend), PE-conjugated anti-CD31 (1:500, BioLegend), PE-conjugated anti-CD140a (1:500, BioLegend) and APC-Cy7-conjugated anti-EpCam (clone G8.8, 1:250, Biolegend), during 45 min at 4 °C on a rocking plate. Living epithelial cells were selected by forward scatter, side scatter, doublets discrimination and by Hoechst dye exclusion. EpCam+/Lin-cells were selected based on the expression of EpCam and the exclusion of CD45, CD31, CD140a (Lin-). Fluorescence-activated cell sorting analysis was performed using FACSAria III and FACS-Diva software (BD Biosciences).

### Histology and immunostaining

For immunostaining on frozen sections, tissues were dissected and embedded in O.C.T. (Tissue Tek) and flash frozen for cryopreservation. For the following staining: EpCam, Sox9, Krt7, tissues were pre-fixed in 4% formaldehyde during 2 h at room temperature, washed in PBS and incubated overnight in a 30% sucrose solution in PBS at 4°C, embedded in O.C.T. (Tissue Tek) and flash frozen for cryopreservation.

For murine and human organoids immunostaining, cells were harvested by disrupting the Matrigel with cold PBS. Organoids were centrifugated, resuspended in 4% formaldehyde for 10 min at room temperature, centrifugated, incubated overnight at 4°C in a 30% sucrose solution in PBS, embedded in O.C.T. and flash frozen for cryopreservation.

Samples were sectioned at 6 µm sections using a M1860 cryostat (Leica Microsystems GmbH). Nonspecific antibody binding was blocked with 5% horse serum (HS), 1% Bovine Serum Albumin (BSA), and 0.2% Triton X-100 during 1 h at room temperature.

For phospho-Smad2 staining on tissue, freshly cut slides were first incubated in 4% formaldehyde during 30min at 4°C before the blocking. Primary antibodies were incubated overnight at 4°C in blocking buffer. Sections were rinsed three times in PBS and incubated with secondary antibodies during 1 h at room temperature. Nuclei were stained with Hoechst (4 mM). Slides were mounted using Glycergel (Dako) supplemented with 2.5% DABCO (Sigma-Aldrich). The following primary antibodies were used: anti-Krt14 (polyclonal chicken, 1:10000, Biolegend, RRID:AB_2616962), anti-GFP (polyclonal goat, 1:1000, Abcam, RRID:AB_305643), anti-p63 (rabbit monoclonal, 1:1000, Abcam, RRI-D:AB_10971840), anti-EpCam (rat polyclonal, 1:500, Biolegend, RRID:AB_1089027), anti-Krt7 (rabbit monoclonal, 1:1000, Abcam, RRID:AB_2783822), anti-Sox9 (rabbit polyclonal, 1:10000, Merck, RRID:AB_2239761), anti-Krt13 (rabbit monoclonal, Abcam, RRID:AB_2134681), anti-Sox2 (rabbit monoclonal, Abcam, RRID:AB_10585428), anti-E-cadherin (rat monoclonal, Thermo Fisher Scientific, RRID:AB_86571), anti-Tenascin C (rat monoclonal, Thermo Fisher Scientific, RRID:AB_2256026), anti-a6 integrin (rat monoclonal, BioLegend,RRID:AB_345296), anti-phospho-Smad2 (rabbit monoclonal, Cell Signaling Technology, RRID:AB_2798798).

The following secondary antibodies were used: anti-rabbit, anti-goat, anti-rat, anti-chicken, conjugated to AlexaFluor488 (1:500, Jackson ImmunoResearch), to rhodamine Red-X (1:500, Jackson ImmunoResearch) or to Cy5 (1:1000, Jackson ImmunoResearch).

### Image acquisition and quantification

All samples from the same experiment were imaged with the same exposure settings. Imaging was performed on a Zeiss Axio Imager M2 fluorescence microscope with a Zeiss Axiocam 503 mono camera for immunofluorescence microscopy using Zen Blue (Zeiss) software. Subsequent quantifications were performed with QuPath software.

### Western blot

Murine WT organoids were treated for 24 hours. After Matrigel disruption, organoids were centrifugated, washed three times with PBS and lysed with fresh RIPA buffer (150 mM NaCl, 0.1% Triton X-100, 0.5% Sodium deoxycholate, 0.1% SDS, 50 mM TrisHCl pH8) supplemented with 1:100 Protease/Phosphatase Inhibitor Cocktail (Cell Signaling Technology, #5872). After 20 min incubation at 4°C, lysates were cleared by centrifugation at 14,000 x g for 10min at 4°C and normalized based on total protein content measured using the Bradford assay (Bio-Rad). Equal protein amounts were resolved on sodium dodecyl sulfate polyacrylamide gel electrophoresis (SDS-PAGE) and transferred into nitrocellulose membranes that were blocked with 5% BSA diluted in TBS 0.1% Tween solution. Proteins were detected with the corresponding primary antibodies. Secondary goat anti-rabbit antibodies (Cell Signaling Technologies, #7074) conjugated to horseradish peroxidase were used to reveal the proteins. Immunoblot analysis was conducted using enhanced chemiluminescence (ECL) reagent according to manufacturer’s protocol (PerkinElmer, Brussels, Belgium) and Images were obtained on Azure c600 (Azure Biosystems, Sierra Court, CA, USA). The following primary antibodies were used: anti-Smad2 (rabbit monoclonal, Cell Signaling Technology, #5339), anti-Smad1 (rabbit monoclonal, Cell Signaling Technology, #12534), anti-Smad5 (rabbit monoclonal, Cell Signaling Technology, RRID:AB_2797946), anti-GAPDH (rabbit monoclonal, Cell Signaling Technology, RRID:AB_561053), anti-phospho-Smad2 (rabbit monoclonal, Cell Signaling Technology, #3108), anti-phospho-Smad1/5 (rabbit monoclonal, Cell Signaling Technology, #9516).

### RNA extraction and quantitative real-time PCR

Organoids and FACS-sorted cells were collected into TRK lysis buffer (Omega bio-tek) and RNA was extracted using E.Z.N.A Total RNA Kit (Omega bio-tek) according to the manufacturer’s recommendations with DNase I digestion protocol on column (Omega bio-tek). After nanodrop RNA quantification, the first strand cDNA was synthesized, using Superscript II (Invitrogen) and random hexamers (Roche) in 25 ul final volume. Quantitative PCR assays were performed using 2 ng of cDNA as template, PowerUP SYBRGreen master mix (Life Technologies Limited) and a Quantstudio 3 real-time PCR system (Applied Biosystems). Actin beta housekeeping gene was used for normalization. Murine and human primers were designed using the NCBI primer designing tool – Primer-blast (https://www.ncbi.nlm.nih.gov/tools/primer-blast/) and are presented in table S8 and S9. . Quantitative PCR Analysis was performed by using the ddCt method with Actin beta as a reference for murine samples and with GAPDH for human samples. The entire procedure was repeated in at least three biologically independent samples and always with technical replicates

### RNA-seq and analysis of bulk samples

RNA quality was checked using a Bioanalyzer 2100 (Agilent technologies). Indexed cDNA libraries were obtained using the Ovation Solo RNA-Seq System (NuGen) following manufacturer’s recommendations. The multiplexed libraries were loaded on a NovaSeq 6000 (Illumina) using a S2 flow cell and sequences were produced using a 200 Cycle Kit. Paired-end reads were mapped against the mouse reference genome GRCm38 using STAR software (version 2.5.3a) to generate read alignments for each sample. Annotations Mus_musculus.GRCm38.90.gtf were obtained from ftp.Ensembl.org. After transcripts assembling, gene level counts were obtained using HTSeq ^53^.

Total raw counts were used for subsequent analyzes with degust 4.1.1 ^54^. All analyses were performed using EdgeR, TMM normalization and ‘‘Min gene read count’’ set at 10.

Bulk RNA seq data from control (CTRL, *n* = 6), K5:Smo EpHi 3 weeks (*n* = 4), 7 weeks (*n* = 4) and 12 weeks (*n* = 6) after TAM administration as well as keratinocytes from the squamous junction (*n* = 3) were downloaded under GEO access GSE148874 and processed as described by the authors ^6^. Heatmap was generated using “heatmap.2” function in R version 4.2.1. It represents values in LogCPM, scaled by row for selected genes associated to BMP pathway activation. Those genes were chosen by comparing the list of upregulated genes from K5:Smo 12 weeks after TAM injection with the geneset (GO_RESPONSE_TO_BMP) from GO terms. Common unique genes were selected with Venny 2.1.

Murine organoids derived from wild-type mice treated for 3 days with different conditions were sequenced : (1) TGF-β inhibitor (CTRL), (2) TGF-β inhibitor + BMP4 (BMP4), (3) TGF-β1, and (4) a combination of TGF-β1 + BMP4. Each condition included two independent cultures. The data are available under the GEO accession number GSE291094. MDS plots were generated using Degust 4.1.1 ^54^.For ibuprofen experiments, EpHi cells from K5:Smo mice, either ibuprofen-treated (n = 3) or untreated (*n* = 3) 12 weeks post-TAM administration, were FACS-sorted and sequenced. Data are available under the GEO accession number GSE291095.Venn diagrams showed in figure 4G and supplementary figure 4D were obtained by comparing the top 2000 significantly (*FDR* <0.05) deregulated transcripts in the different conditions. Venn diagrams showed in figure 6H were obtained by comparing the top 1500 deregulated transcripts in the different conditions (with FDR < 0.05 for TGF-β1 + BMP4 vs CTRL and FDR < 0.1 for TGF-β1 + BMP4 + Ibu vs TGF-β1 + BMP4).

### GSEA analysis

Gene set enrichment analysis (GSEA) was performed using the fgsea package ^55^ in R version 4.2.1. Genes were ranked in descending order and then filtered to keep only significant differentially expressed genes: (1) for K5Smo 12w: abs(*LFC*)>1 & *FDR*<0.05 and (2) for murine organoids FDR<0.05. The ranking metric was defined as the values of LFC. The C5 collection adapted for mouse which contains gene sets annotated by GO terms have been downloaded (http://bioinf.wehi.edu.au/software/MSigDB/) and the unbiased gene ontology analyses has been realized with the function “fgseaMultilevel”. Pathways with adjusted *P*-value <0.001 were considered as significant. The “plotEnrichment” function was used for enrichment plot. Dotplot was generated using “ggplot” function in R version 4.2.1.

### ATAC-sequencing and analysis

Data under GEO accession number GSE148872 were visualized using IGV genome browser and the ATAC seq profile of the *Sox9* locus in WT mouse esophagus epithelial cells was exported as JPEG file. FIMO (Find Individual Motif Occurences) was used to identify putative binding motif in the Sox9 promoter-associated peak. Dotplot was generated using “ggplot” function in R version 4.2.1.

### Single cell-RNA sequencing and analysis

Sorted cells were collected in 1 mL of PBS + 0.04% BSA at 4°C. Samples were loaded on a chromium chip. Samples were then processed using the Chromium Single Cell 3ʹ Reagent Kits v3 (10x Genomics) following the manufacturer’s recommendations. The multiplexed libraries were loaded on a NovaSeq 6000 (Illumina) using a S1 flow cell, and paired-end sequences were produced using a 100-cycle kit (Read1, 28 cycles; i7 Index, 8 cycles; i5 Index, 0 cycles; and Read2, 87 cycles). Cell Ranger version 3.1.0 pipeline with the RNA-seq aligner STAR were used to generate output files aligned on mm10 reference genome. Package Seurat v4.2 from Bioconductor (48) was used in R to perform all the analyses.

The analyses followed recommendations from Satija Lab (https://satijalab.org). The datasets were converted into Seurat objects and SoupX v1.6.2 ^56^ was used with default parameters to remove ambient RNA contamination. For each sample, poor quality cells, potential empty droplets and possible multiplets were filtered out based on high content of mitochondrial genes and on number of features. The data were normalized using “SCTransform” that outperforms traditional global scaling normalization methods derived from bulk RNA-seq. To avoid the influence of mitochondrial reads in the analyses, we used “vars.to.regress=percent.mt” with “percent.mt” calculated using “PercentageFeaturesSet.” For each Seurat object, the following steps have been run: PCAs were calculated using “RunPCA”, and Uniform Manifold Approximation and Projection (UMAP) analyses have been done using “RunUMAP”. Distances between cells were defined using “FindNeighbors”. The cells were grouped using “FindClusters” with a resolution of 0.3. Doublets could then be removed using DoubletFinder R package v2.0.3 ^57^. Last, for visualization and differential expression (DE) analysis, RNA counts were normalized using “NormalizeData.” In this study, three samples were sequenced at single-cell level:

1. Control esophagus: dataset was downloaded under GEO access GSE148875 and sample was processed as described by the authors ^6^. A total of 10,672 of these cells were sequenced. The thresholds were defined as follows: nFeature_RNA > 250 and nFeature_RNA < 6000 and percent.mt < 10. DoubletFinder identified 674 doublets that were removed from the dataset. A total of 8,526 cells was used for the analysis. This sample has been used in Figs. 1, 2, 3, and 5 and related supplementary figures.
2. K5:Smo 12 weeks post-TAM esophagus : For all living cells of thoracic esophagus, 110,000 living cells were sorted from a pool of three K5:Smo mice. A total of 20,104 of these cells were sequenced. The thresholds were defined as follows: nFeature_RNA > 250 and nFeature_RNA < 5800 and percent.mt < 15. DoubletFinder identified 2499 doublets that were removed from the dataset. A total of 15,624 cells was used for the analysis. This sample has been used in Figs. 1, 2, 3, and 5 and related supplementary figures. Data are available under GEO accession number GSE291090.
3. K5:Smo 12 weeks post-TAM administration treated with ibuprofen esophagus: For all living cells of thoracic esophagus, 110,000 living cells were sorted from a pool of two ibuprofen-treated k5:Smo mice. A total of 19,925 of these cells were sequenced. The thresholds were defined as follows: nFeature_RNA > 250 and nFeature_RNA < 5000 and percent.mt < 15. DoubletFinder identified 2521 doublets that were removed from the dataset. A total of 15,857 cells was used for the analysis. This sample has been used in Fig.5 and related supplementary figure. Data are available under GEO accession number GSE291090.

The three samples were merged using the “merge” function (with the parameter *merge.data = TRUE*). Cluster annotation was performed based on differentially expressed markers for each group. For epithelial and immune cell annotation, subsetting was performed based on *Epcam* and the co-expression of *Vim* and *Ptprc* respectively. After running SCT normalization, “RunPCA”, “FindNeighbors”, and “FindClusters”, cells could be accurately annotated. In the whole dataset, clusters containing less than 20 cells were removed. Dot plots and violin plots were generated using “DotPlot” and “VlnPlot” function respectively.

Characterization of the signaling pathway network was performed using *CellChat* ^7^, following the authors’ recommendations. Bar plots comparing the overall information flow of each signaling pathway were generated using the "rankNet" function, with the argument *target.use* defining epithelial cell clusters in the control sample and EpHi epithelial clusters in the K5:Smo samples, and a *P*-value cutoff of 0.01. Circle plots were generated using “netVisual_aggregate” function.

### Quantification and statistical analysis

Statistical and graphical data analyses were performed using Prism 8 (GraphPad) and R software. All experiments presented were replicated at least twice. Data in histograms represent the mean unless otherwise specified in the legend. For qPCR analysis, statistical significance was assessed using the Kruskal-Wallis test followed by Dunn’s multiple comparisons test, unless stated otherwise in the legend. For histological quantification, statistical significance was assessed using the ANOVA test followed by Tukeys’s multiple comparisons test. *P*-value < 0.05 was considered statistically significant. All tests are two-sided.

## Resource availability

### Lead Contact

Further information and requests for resources and reagents should be directed to and will be fulfilled by the Lead Contact, Benjamin Beck.

### Materials availability

This study did not generate new unique reagents.

### Data Availability

The dataset from RNA-sequencing and single-cell RNA-sequencing generated during this study will be available at Gene Expression Omnibus under the following accession number: GSE291094 (RNA-seq data from WT organoids), GSE291095 (RNA-seq data from ibuprofen-treated K5:Smo mice), GSE291090 (single-cell RNA-sequencing).

