## Supplementary figures 1 to 7 for "Sox9-dependent plasticity of esophageal progenitors is fine-tuned by cues from the microenvironment"

### Supplementary Figure1

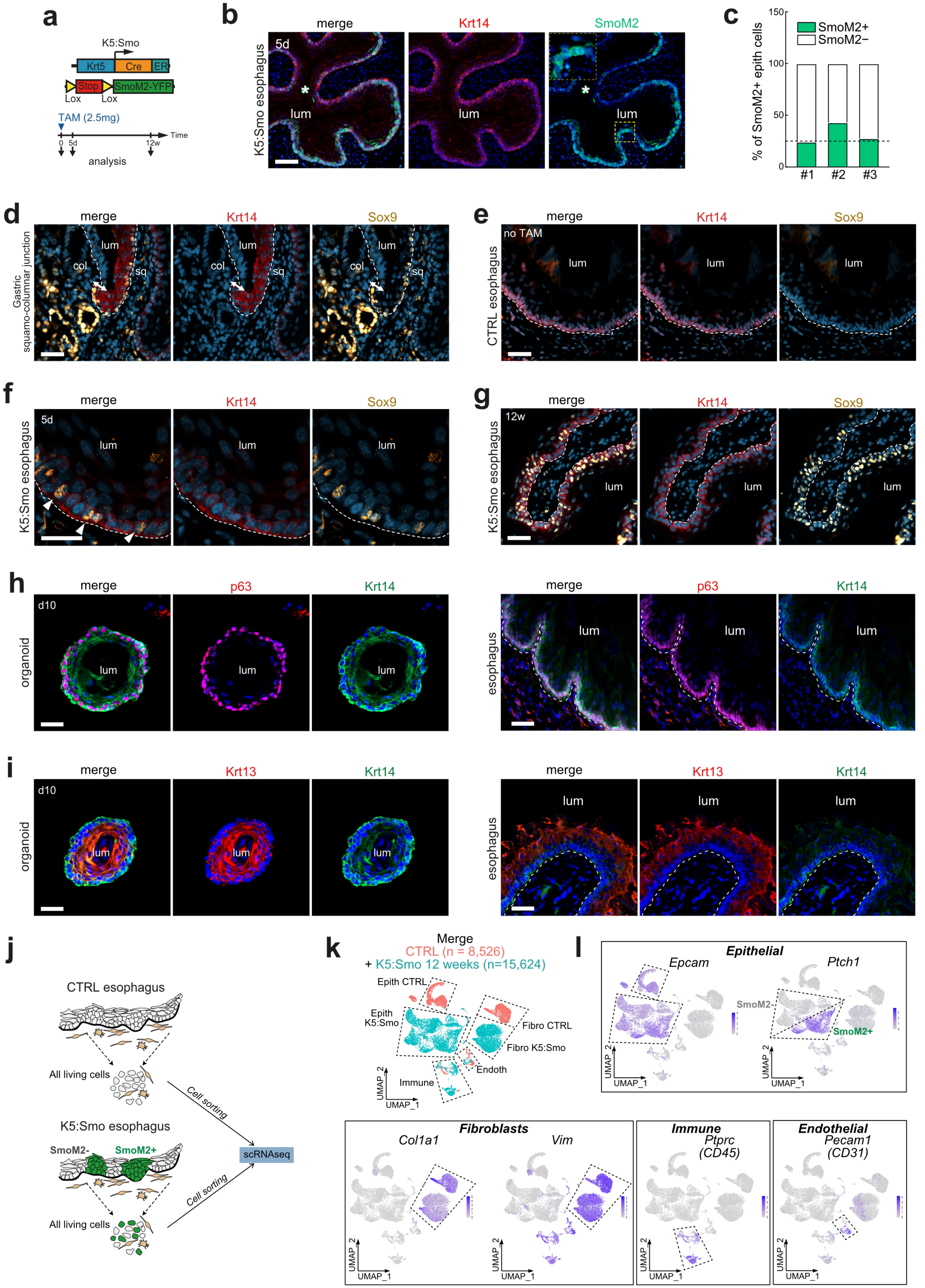

**Supplementary Fig. 1. | Sox9 expression at the squamo-columnar junction and in esophageal progenitors**

(a) Experimental design
(b) Co-immunostaining for Krt14 and SmoM2 (YFP) in K5:Smo esophagus 5 days after TAM. Scale bars represent 100 µm.
(c) Quantification of the proportion of SmoM2 epithelial cells 5 days after TAM (*n* = 3 mice).
(d) Co-immunostaining for Krt14 and Sox9 at the SCJ of CTRL mice.
(e) Co-immunostaining for Krt14 and Sox9 in CTRL esophagi (no TAM).
(f) Co-immunostaining for Krt14 and Sox9 in K5:Smo esophagi 5 days post-TAM. Scale bar represents 20 µm.
(g) Co-immunostaining for Krt14 and Sox9 in K5:Smo esophagi 12 weeks post-TAM.
(h) Co-immunostaining for Krt14 and p63 in esophageal organoids grown for 13 days and in the esophagus of CTRL mice.
(i) Co-immunostaining for Krt13 and Krt14 in esophageal organoids grown for 13 days and in the esophagus of CTRL mice.
(j) Experimental design to conduct scRNA seq study in CTRL and K5:Smo 12 weeks post-TAM esophagi. All living cells were FACS-sorted and sequenced.
(k) UMAP visualization merged scRNA seq data from all living esophageal cells from CTRL mice (*n* = 8,526 cells) and K5:Smo (*n* = 15,624 cells) 12 weeks after TAM mice.
(l) UMAP visualization of gene expression patterns using classical markers associated with epithelial, fibroblastic, endothelial and immune lineages. The clusters of immune and epithelial cells were further subsetted for better clarity. Subset and annotation are presented in Fig. S2B.

Merge channels of the IF are represented in the Fig. 1.
Nuclear staining is represented in blue. Scale bars represent 50 µm. TAM, tamoxifen administration; CTRL, control; Lum, lumen; Col, columnar; Sq, squamous; SCJ, squamo-columnar junction; dash lines represent the basal lamina.

### Supplementary Figure2

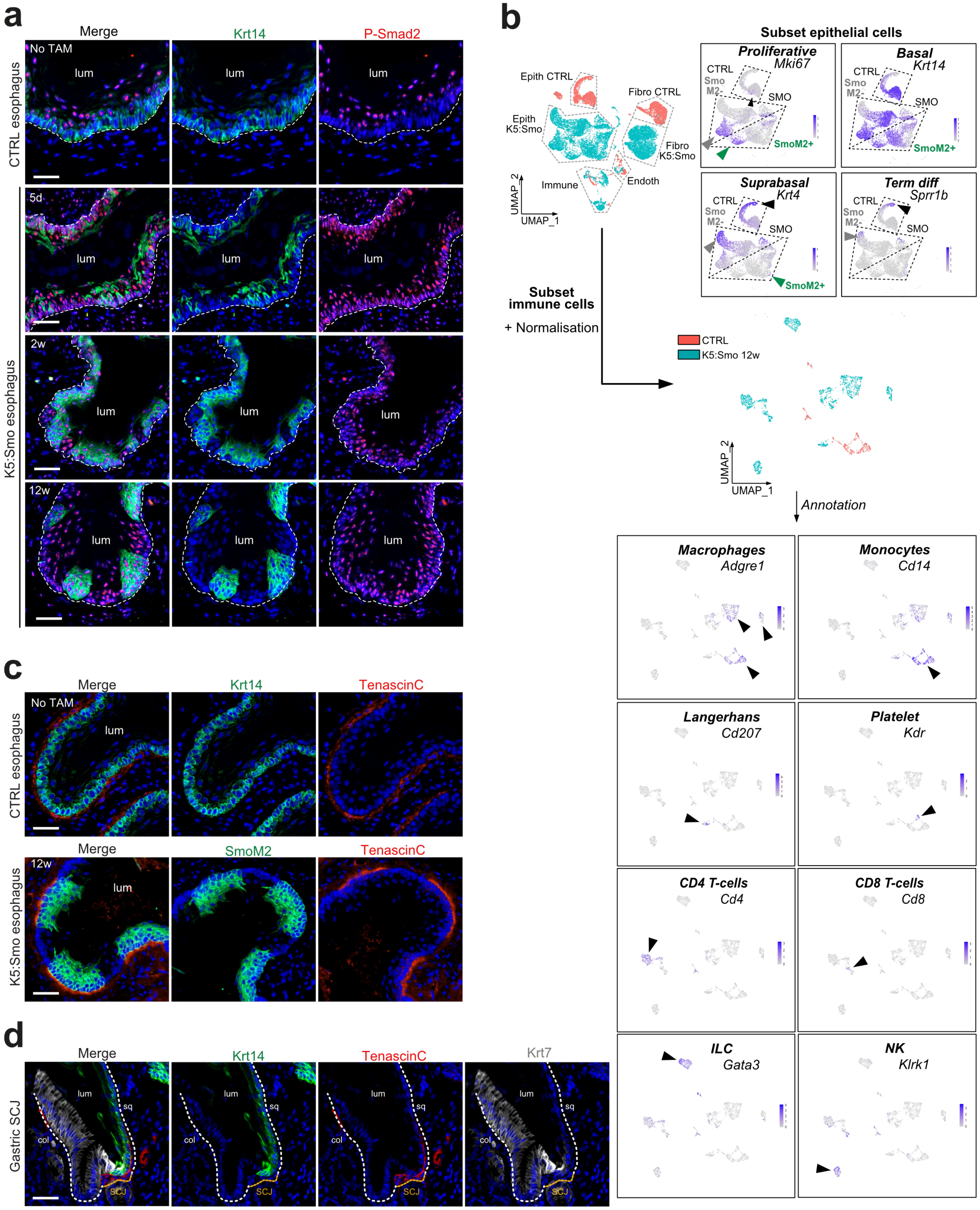

**Supplementary Fig. 2. | Hedgehog pathway stimulates TGF-β1/Smad2 pathway activation in esophageal cells**

(a) Co-immunostaining for Krt14 and P-Smad2 in CTRL esophagus and SmoM2 (YFP) and P-Smad2 in K5:Smo esophagus 5 days, 2weeks and 12 weeks after TAM.
(b) UMAP visualization of gene expression patterns using classical markers associated with the different epithelial and immune populations. These data were used for the annotation of clusters represented in Fig. 2C, 2D, 3D, 3E and 5A.
(c) Co-immunostaining for Krt14 and Tenascin-C in CTRL esophagi; SmoM2 (YFP) and Tenascin-C in K5:Smo esophagi 12 weeks post TAM.
(d) Co-immunostaining for Krt14, Tenascin-C and Krt7 in the gastric SCJ of CTRL mice.

Merge channels of the IF are represented in the Fig. 2.
Nuclear staining is represented in blue. Scale bars represent 50 µm. TAM, tamoxifen administration; col, columnar; sq, squamous; SCJ, squamo-columnar junction; lum, lumen; dash lines represent the basal lamina.

### Supplementary Figure3

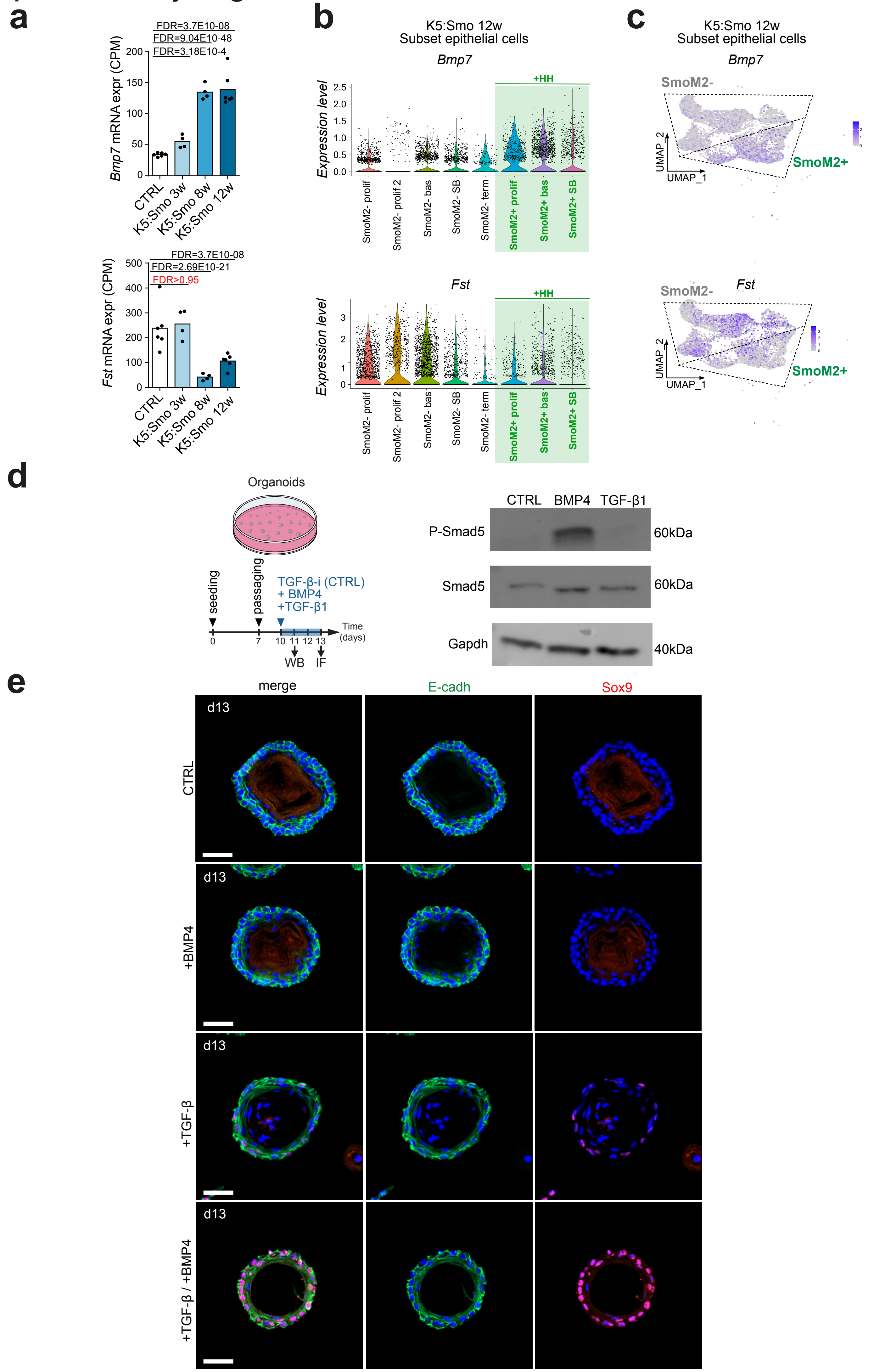

**Supplementary Fig. 3. | Hedgehog pathway activation in esophageal progenitors progressively induces a Bmp7 autocrine loop**

(a) *Bmp7 and Fst* mRNA expression measured by RNA seq in FACS-sorted K5:Smo SmoM2+ esophageal cells (3 and 12 weeks post-TAM) and in CTRL condition.

(b) Violin plots showing Bmp7 and *Fst* expression in scRNA-seq data from K5:Smo mice, 12 weeks post-TAM. *Bmp7* is upregulated in K5:Smo SmoM2+ cells compared to SmoM2- neighboring epithelial cell. *Fst* is downregulated in K5:Smo SmoM2+ cells compared to SmoM2- neighboring epithelial cells.

(c) UMAP visualization of *Bmp7* and *Fst* expression in scRNA seq data from K5:Smo mice, 12 weeks post-TAM.

(d) (Left) Experimental design. (Right) Expression of P-Smad1/5 and Smad1/5 measured by western blot in esophageal organoids treated for 24 hours with TGF-β inhibitor, BMP4 or TGF-β1. This data shows that Smad1/5 is phosphorylated only following BMP pathway activation *in vitro*.

(e) Co-immunostaining of E-cadherin and Sox9 in esophageal organoids treated for 3 days with TGF-β inhibitor (CTRL), BMP4, TGF-β1 or TGF-β1 + BMP4.

Merge channels of the IF are represented in the Fig. 3.
Nuclear staining is represented in blue. Scale bars represent 50 µm. TAM, tamoxifen administration.

### Figure S4

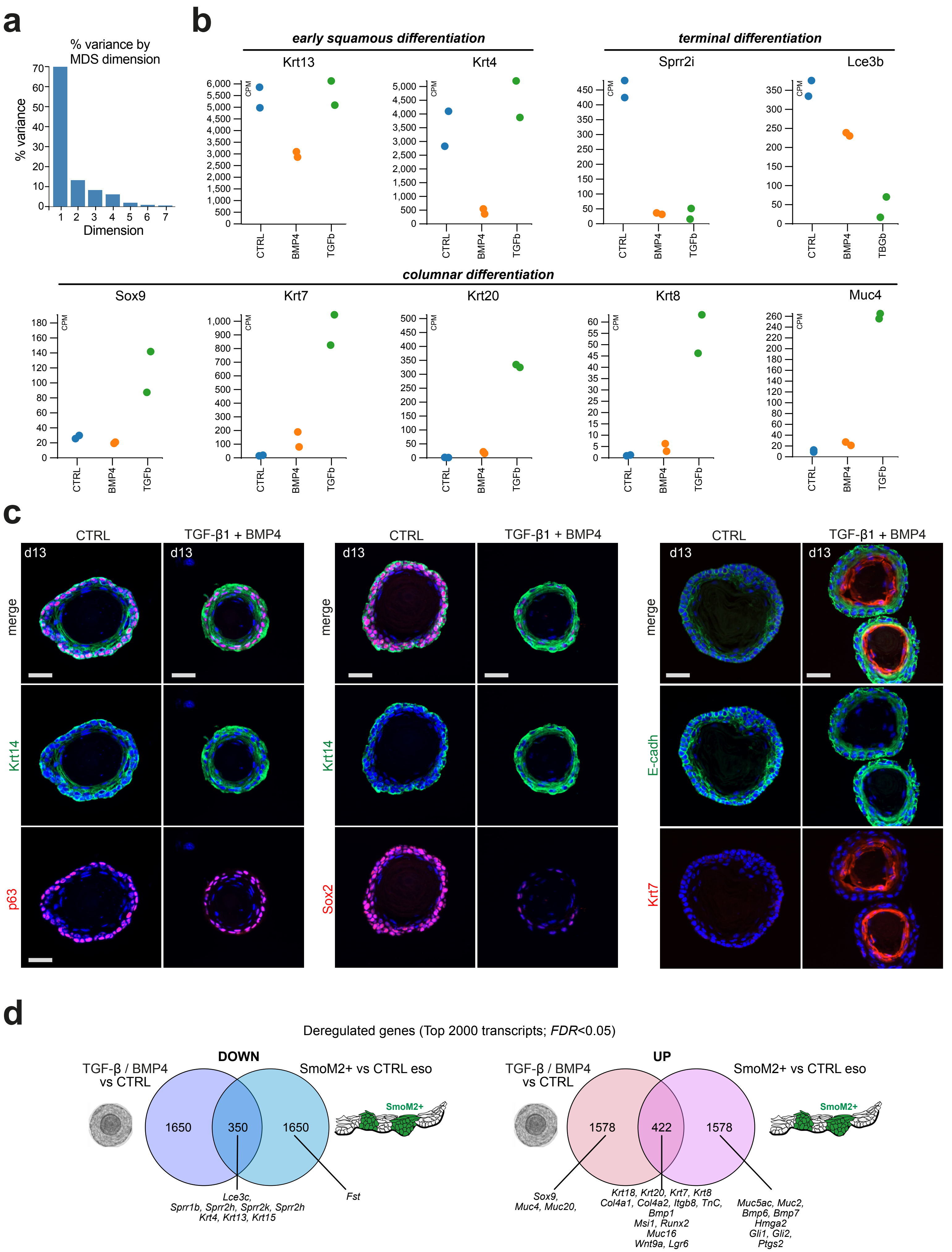
a
% variance by MDS dimension
% variance
Dimension
b
early squamous differentiation
terminal differentiation
Krt13
Krt4
Sprr2i
Lce3b
columnar differentiation
Sox9
Krt7
Krt20
Krt8
Muc4
CPM
CTRL
BMP4
TGFb
TBGb
c
CTRL
TGF-β1 + BMP4
d13
merge
Krt14
p63
Sox2
E-cadh
Krt7
d
Deregulated genes (Top 2000 transcripts; FDR<0.05)
DOWN
TGF-β / BMP4 vs CTRL
SmoM2+ vs CTRL eso
1650
350
1650
SmoM2+
Lce3c,
Sprr1b, Sprr2h, Sprr2k, Sprr2h
Krt4, Krt13, Krt15
Fst
UP
TGF-β / BMP4 vs CTRL
SmoM2+ vs CTRL eso
1578
422
1578
SmoM2+
Sox9,
Muc4, Muc20,
Krt18, Krt20, Krt7, Krt8
Col4a1, Col4a2, Itgb8, TnC,
Bmp1
Msi1, Runx2
Muc16
Wnt9a, Lgr6
Muc5ac, Muc2,
Bmp6, Bmp7
Hmga2
Gli1, Gli2,
Ptgs2

**Supplementary Fig. 4. | TGF-β/BMP signaling pathway activation in esophageal progenitors *in vitro* partially recapitulates the phenotype of HH-stimulated cells *in vivo***

(a) Barplot representing the percentage of variance of the MDS plot shown in Fig. 4B.
(b) Gene expression plots of squamous (*Krt13*, *Krt4, Sprr2i,* and *Lce3b*) and columnar (*Sox9*, *Krt7*, *Krt20*, *Krt8,* and *Muc4*) markers obtained by RNA seq of esophageal organoids treated with TGF-β inhibitor (CTRL), BMP4 or TGF-β1.
(c) Co-immunostaining of Krt14 and p63; Krt14 and Sox2; E-cadh and Krt7 in esophageal organoids treated with TGF-β inhibitor (CTRL) or TGF-β1 + BMP4.
(d) Venn diagrams comparing the top 2000 significantly deregulated transcripts in (1) organoids treated with TGF-β1 + BMP4 compared to CTRL and (2) in SmoM2-positive keratinocytes of K5:Smo mice 12 weeks post TAM mice compared to normal esophageal keratinocytes. These data show that 17.5% of downregulated genes and 21.1% of upregulated genes are common between the TGF-β/BMP4-treated organoids and SmoM2-positive keratinocytes of K5:Smo mice.

Merge channels of the IF are represented in the Fig. 4.
Nuclear staining is represented in blue. Scale bars represent 50 µm; TAM, tamoxifen administration.

### Supplementary Figure5

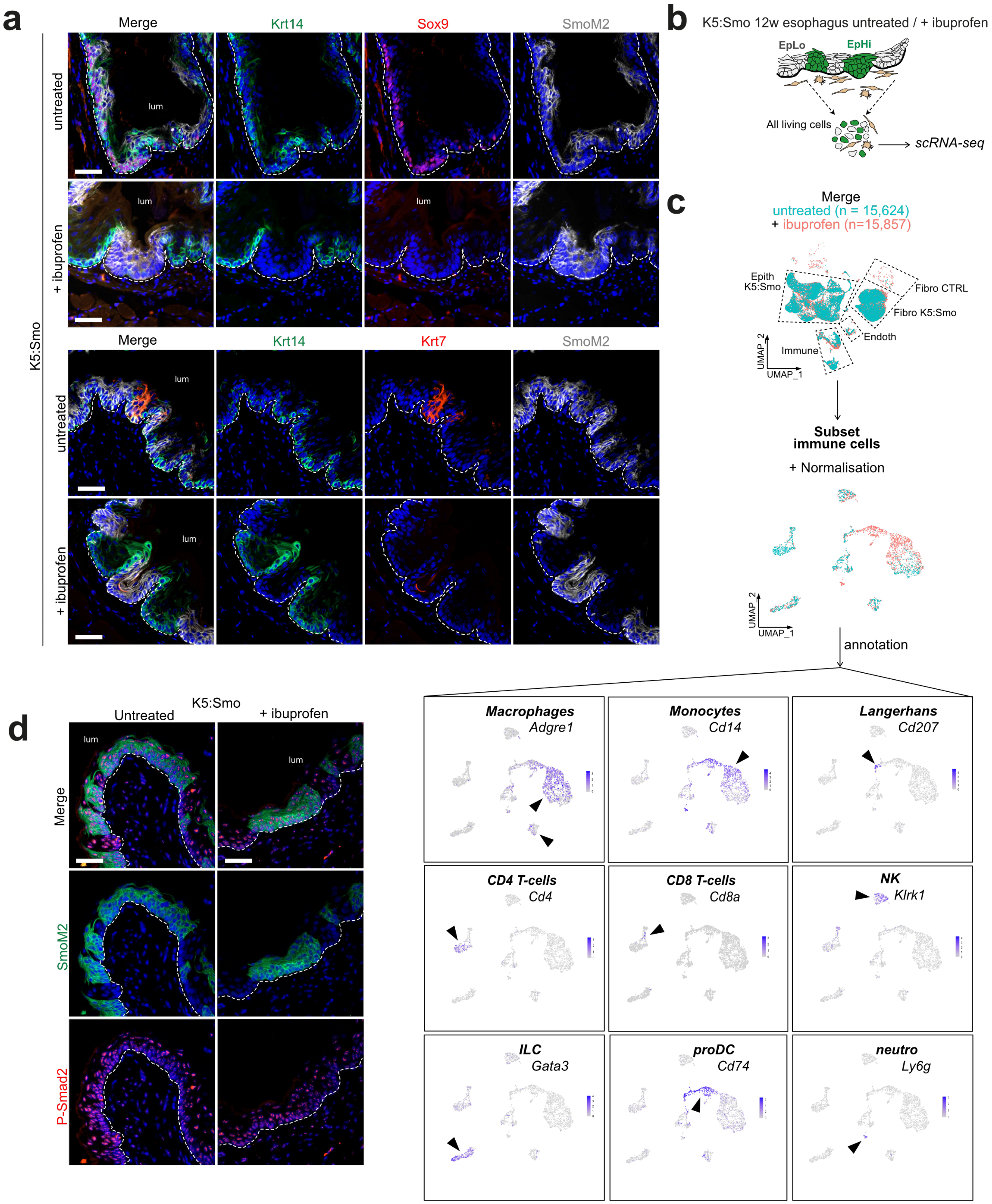

**Supplementary Fig. 5. | Ibuprofen inhibits HH-induced Sox9 expression in esophageal progenitors *in vivo***

(a) Co-immunostaining for Krt14, Sox9, EpCam (surrogate of SmoM2 expression) and Krt14, Krt7, Epcam in untreated and ibuprofen-treated K5:Smo esophagi 12 weeks post-TAM.
(b) Experimental design.
(c) UMAP visualization of merged scRNA-seq data from all living esophageal cells from untreated K5:Smo 12 weeks post-TAM mice (*n* = 15,624 cells) and ibuprofen-treated K5:Smo 12 weeks post-TAM mice (*n* = 15,857 cells). Gene expression patterns using classical markers associated with the different immune populations are represented. These data were used for the annotation of clusters represented in Fig. 5F.
(d) Co-immunostaining for SmoM2-YFP and P-Smad2 in untreated and ibuprofen-treated K5:Smo esophagi 12 weeks post-TAM.

Merge channels of the IF are represented in the Fig. 5.
Nuclear staining is represented in blue. Scale bars represent 50 µm; TAM, tamoxifen administration; dash lines represent the basal lamina.

### Supplementary Figure6

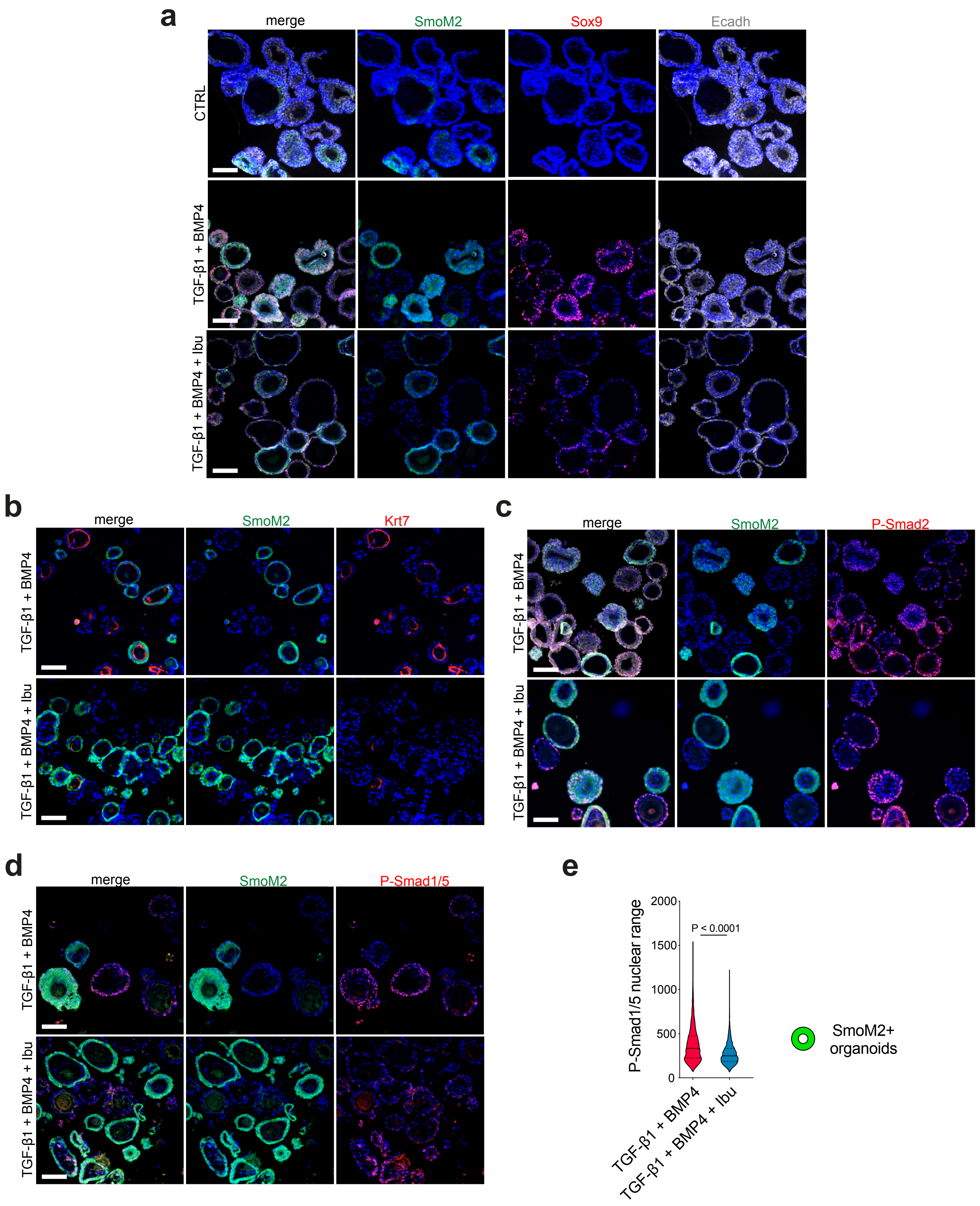

**Supplementary Fig. 6. | Ibuprofen inhibits TGF-β/BMP-stimulated Sox9 protein expression but not its mRNA expression in mouse esophageal progenitors *ex vivo***

(a) Co-immunostaining for SmoM2-YFP, Sox9 and E-cadh in K5:Smo esophageal organoids treated with TGF-β inhibitor (CTRL), TGF-β1 + BMP4 or TGF-β1 + BMP4 + ibuprofen.
(b) Co-immunostaining for SmoM2-YFP and Krt7 in K5:Smo esophageal organoids treated with TGF-β1 + BMP4 or TGF-β1 + BMP4 + ibuprofen.
(c) Co-immunostaining for SmoM2-YFP and P-Smad2 in K5:Smo esophageal organoids treated with TGF-β1 + BMP4 or TGF-β1 + BMP4 + ibuprofen.
(d) Co-immunostaining for SmoM2-YFP and P-Smad1/5 in K5:Smo esophageal organoids treated with TGF-β1 + BMP4 or TGF-β1 + BMP4 + ibuprofen.
(e) Boxplot (min to max) summarizing Smad1/5 protein phosphorylation measured by immunofluorescence in K5:Smo esophageal organoid treated for 3 days with TGF-β1 + BMP4 or TGF-β1 + BMP4 + ibuprofen.

Merge channels of the IF are represented in the Fig. 6.
Nuclear staining is represented in blue. Scale bars represent 100 µm.

### Supplementary Figure7

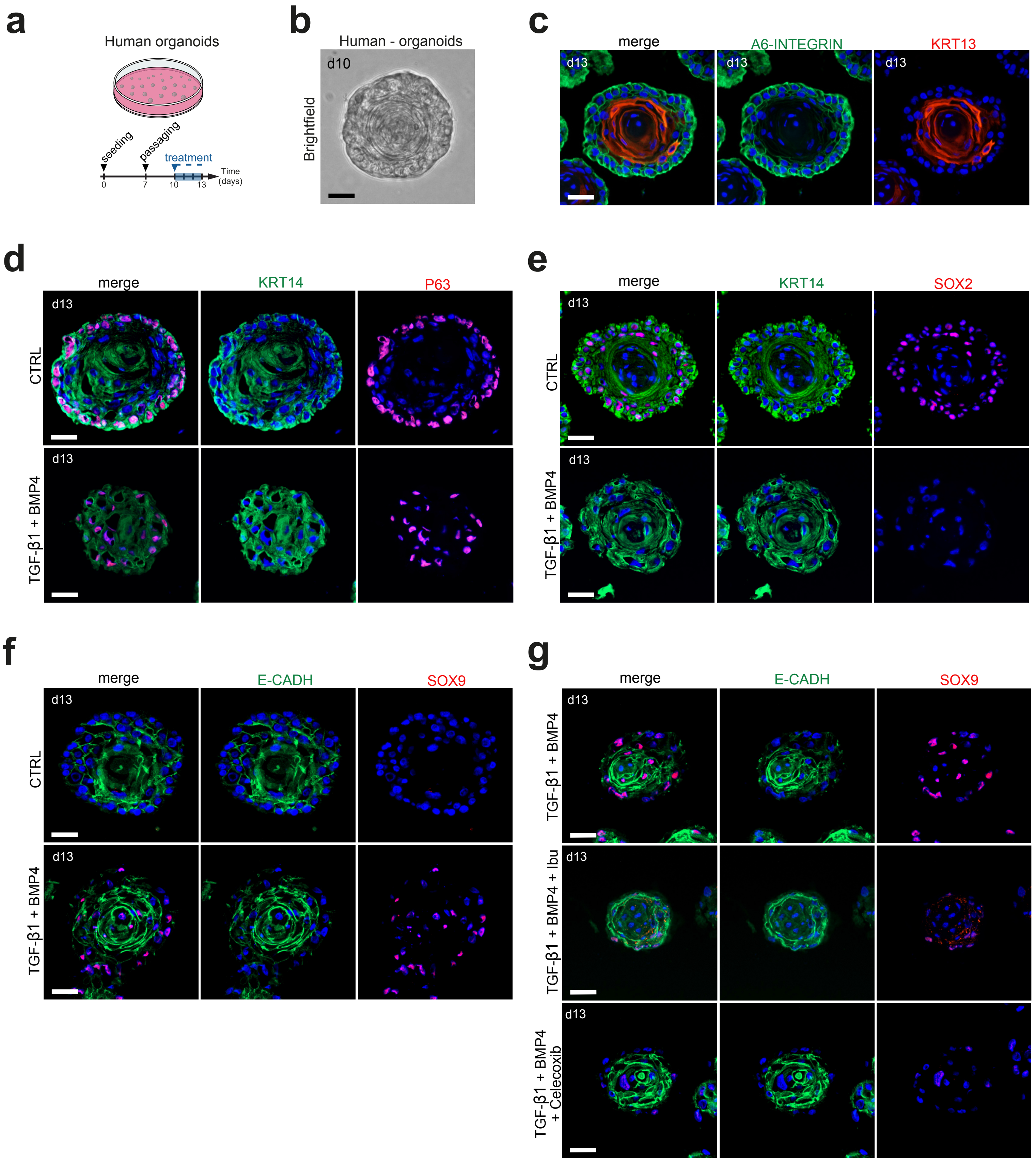

**Supplementary Fig. 7. | Ibuprofen inhibits TGF-β/BMP-stimulated SOX9 protein expression but not its mRNA expression in human esophageal progenitors *ex vivo***

(a) Experimental design.
(b) Brightfield image of human esophageal organoid after 10 days in culture.
(c) Co-immunostaining for A6-INTEGRIN and KRT13 in human esophageal organoids.
(d) Co-immunostaining for KRT14 and P63 in human esophageal organoids treated with TGF-β inhibitor (CTRL) or TGF-β1 + BMP4.
(e) Co-immunostaining for KRT14 and SOX2 in human esophageal organoids treated with TGF-β inhibitor (CTRL) or TGF-β1 + BMP4.
(f) Co-immunostaining for E-CADH and SOX9 in human esophageal organoids treated with TGF-β inhibitor (CTRL) or TGF-β1 + BMP4.
(g) Co-immunostaining for E-CADH and SOX9 in human esophageal organoids treated with TGF-β1 + BMP4, TGF-β1 + BMP4 + ibuprofen or TGF-β1 + BMP4 + celecoxib (a specific COX-2 inhibitor).

Merge channels of the IF are represented in the Fig. 7.
Nuclear staining is represented in blue. Scale bars represent 50 µm.
