## Supplementary Tables 1 to 9 for "Sox9-dependent plasticity of esophageal progenitors is fine-tuned by cues from the microenvironment"

### Table S1 : Putative binding motifs identified in the open chromatin region upstream of *Sox9* using *FIMO* (*qvalue*<0.01)

| motifID | TF | start | stop | strand | *score* | *pval* | *qvalue* | matchedSequence |
|---|---|---|---|---|---|---|---|---|
| MA1721.2 | ZNF93 | 1405 | 1418 | - | 18,50 | 3,5E-07 | 0,000667 | GGCTGCGGCGGCGG |
| MA1961.2 | PATZ1 | 744 | 754 | + | 17,64 | 7,6E-07 | 0,00156 | GGGGGTGGGGG |
| MA1630.3 | ZNF281 | 744 | 753 | + | 17,16 | 1,2E-06 | 0,00283 | GGGGGTGGGG |
| MA1980.1 | ZNF418 | 282 | 296 | + | 16,45 | 1,4E-06 | 0,00416 | AGGAGGCAAGAAGCA |
| MA0816.1 | Ascl2 | 671 | 680 | - | 16,20 | 2,3E-06 | 0,0059 | AACAGCTGCT |
| MA0795.1 | SMAD3 | 770 | 779 | + | 16,07 | 3,1E-06 | 0,00704 | AGTCTAGACA |
| MA0595.1 | SREBF1 | 970 | 979 | + | 15,85 | 2,9E-06 | 0,00749 | CTCACCCCAC |
| MA0747.2 | SP8 | 741 | 751 | - | 15,75 | 1,9E-06 | 0,00523 | CCACCCCCACT |
| MA0741.1 | KLF16 | 1522 | 1532 | + | 15,53 | 3,4E-06 | 0,00753 | CACCCGCCCCC |
| MA1540.3 | NR5A1 | 366 | 377 | + | 15,52 | 2,9E-06 | 0,00851 | AGTTCAAGGTCG |
| MA1627.2 | Wt1 | 2 | 11 | + | 15,43 | 3,8E-06 | 0,00908 | CCTCCCCCAG |
| MA1557.1 | SMAD5 | 770 | 779 | + | 15,25 | 4,0E-06 | 0,00675 | AGTCTAGACA |
| MA0073.2 | RREB1 | 748 | 766 | - | 14,49 | 1,6E-06 | 0,00203 | ACCCTCCACAACCCCCCAC |
| MA1976.2 | ZNF320 | 941 | 960 | - | 14,19 | 3,9E-06 | 0,00699 | AGGGGGGCGCGGGGCAAGCG |
| MA1107.3 | KLF9 | 742 | 752 | - | 14,02 | 8,6E-06 | 0,00271 | CCCACCCCCAC |
| MA1596.1 | ZNF460 | 1426 | 1441 | + | 12,84 | 4,6E-06 | 0,00915 | GCGCCAACCTCCCCGG |
| MA1155.1 | ZSCAN4 | 794 | 808 | + | 7,31 | 1,9E-05 | 0,00596 | CGCACACACACACAC |

### Table S2 : Tissue digestion

| Reagent or resource | Source | Identifier |
|---|---|---|
| Collagenase I | A.G. Scientific | Cat # C-2823 |
| Foetal Bovine Serum | Gibco | Cat# 10270-106 |
| Trypsin solution 2.5% | A&E Scientific | Cat # TRY-2B10 |
| Dulbecco's PBS | Sigma-Aldrich® | Cat # D8537 |

### Table S3: FACS and flow cytometry antibodies

| Reagent or resource | Source | Identifier |
|---|---|---|
| Rat anti-CD45 PE | BioLegend | Cat #103106; RRID:AB_312971 |
| Rat anti-CD140a PE | BioLegend | Cat #135906; RRID:AB_1953269 |
| Rat anti-CD31 PE | BioLegend | Cat #102508; RRID:AB_312915 |
| Rat anti-CD326 (EpCam) APC-Cy7 | BioLegend | Cat #118218; RRID:AB_2098648 |

**Table S4 : Immunostaining & Western Blot antibodies**

| Reagent or resource | Species, dilution | Source | Identifier |
|---|---|---|---|
| anti-Krt14 | Chicken, 1:30000 | Biolegend | Cat #906004; RRID_AB_2616962 |
| anti-GFP | Goat, 1:1000 | Abcam | Cat # ab6673; RRID_AB_305643 |
| anti-p63 | Rabbit, 1:1000 | Abcam | Cat # ab124762; RRID_AB_10971840 |
| anti-EpCam | Rat, 1:1000 | Biolegend | Cat #118202; RRID_AB_1089027 |
| anti-Krt7 | Rabbit, 1:1000 | Abcam | Cat # ab181598; EPR17078 |
| anti-Sox9 | Rabbit, 1:10000 | Merck | Cat # AB5535; RRID_AB_2239761 |
| anti-P-Smad2 | Rabbit, 1:75 | Cell Signaling Technology | Cat #18338 |
| anti-E-cadherin | Rat 1:500 | Invitrogen | Cat #13-1900; RRID_ AB_2533005 |
| anti-Itga6 | Rat 1:200 | Biolegend | Cat #313 602; RRID_AB_345296 |
| anti-Krt13 | Rabbit 1:1000 | Abcam | Cat # ab92551; EPR3671 |
| anti-Tnc | Rat 1:1000 | Invitrogen | Cat # MA1-26778; RRID_AB_2256026 |
| anti-Sox2 | Rabbit 1:200 | Abcam | Cat # Ab92494 RRID:AB_10585428 |
| anti-Gapdh (WB) | Rabbit 1:1000 | Cell Signaling Technology | Cat #2118 |
| anti-Smad2 (WB) | Rabbit 1:1000 | Cell Signaling Technology | Cat #5339 |
| anti-P-Smad2 (Ser465/467) (WB) | Rabbit 1:500 | Cell Signaling Technology | Cat #3108 |
| Smad1 | Rabbit 1:1000 | Cell Signaling Technology | Cat #6944 |
| Smad5 | Rabbit 1:1000 | Cell Signaling Technology | Cat #12534 |
| P-Smad1/5 | Rabbit 1:1000 (western blot) and 1:100 (immunostaining) | Cell Signaling Technology | Cat #9516 |

**Table S5: Murine/human KSFM medium**

| Reagent or resource | Concentration | Source | Identifier |
|---|---|---|---|
| KSFM medium | N.A | Thermofisher | Cat #10725018 |
| Bovine Pituitary Ex-tract (BPE) | 50 µg/ml | Thermofisher | Cat #37000015 |
| Penicillin/Streptomycin (100x) | 20000U | Thermofisher | Cat #15070063 |
| Human Recombinant Epidermal Growth Fac-tor (EGF1-53) | 1 ng/ml | Thermofisher | Cat #37000015 |
| $CaCl_2.2H_2O$ | 0.5 mM | Sigma-Aldrich® | Cat #223506-25G |
| Fungizone | 250 ng/ml | Gibco | Cat #15290026 |

**Table S6: Culture reagents**

| Reagent or resource | Source | Identifier |
|---|---|---|
| Matrigel | Corning® | Cat #354234 |
| Matrigel growth factor reduced | Corning® | Cat #356234 |
| Rock inhibitor (Y-27632, dihydrochloride) | Sigma-Aldrich® | Cat # Y0503 |
| Ibuprofen | Sigma-Aldrich® | Cat # V900344 |
| Shh (high activity) | Biotechne | Cat #908-SH-005 |
| Recombinant mouse TGFβ1 | Biolegend | Cat #763104 |
| Recombinant mouse BMP-4 | Biotechne | Cat #5020-BP-010 |
| A83-01 (TGFβ inhibitor) | Sigma-Aldrich® | Cat # SML 0788 |
| Recombinant human TGFβ1 | Biolegend | Cat #580704 |
| Celecoxib | Selleckchem | Cat # S1261 |

**Table S7: Additional reagents**

| Reagent or resource | Source | Identifier |
|---|---|---|
| Isopentane | VWR | Cat #24872.323 |
| O.C.T | Tissue Tek | Cat #4583 |
| Formaldehyde 4% | VWR | Cat #116 994 55 |
| Sucrose 30% | Homemade:<br>300gr sucrose<br>1L PBS1x<br>Sodium azide | N.A |
| Horse serum | Capricorn Scientific | Cat # HOS-1b |
| Bovine serum albumin | Sigma-Aldrich® | Cat #810531 |
| Triton X-100 | Sigma-Aldrich® | Cat # T8787 |
| Mounting medium Glycergel | Dako | Cat # C0563 |
| DABCO | Sigma-Aldrich® | Cat # D27802-100G |
| Hoechst | Thermofisher | Cat # H3569 |
| RIPA buffer | Homemade:<br>NaCl 150 mM<br>Triton X-100 0.1%<br>Sodium deoxycholate 0.5%<br>SDS 0.1%<br>TRIS-HCl 50 mM pH8 | N.A |
| Tween20 | Bio-Rad | Cat #1706531 |
| Bradford assay | Bio-Rad | Cat #5000006 |
| Protease inhibitor | Cell Signaling Technology | Cat #5872 |

**Table S8: Mouse qPCR primers**

| Gene name | Forward (5' – 3') | Reverse (5' – 3') |
|---|---|---|
| *Bactin* | CACTGTCGAGTCGCGTCC | TCATCCATGGCGAACTGGTG |
| *Gli1* | CCGACGGAGGTCTCTTTGTC | AACATGGCGTCTCAGGGAAG |
| *Sox9* | CACAAGAAAGACCACCCCGA | GGACCCTGAGATTGCCCAGA |
| *Krt7* | CCGGAATGAGATTGCGGAGA | CTCTAACTTGGCACGCTGGT |

**Table S9: Human qPCR primers**

| Gene name | Forward (5' – 3') | Reverse (5' – 3') |
|---|---|---|
| *SOX9* | GCTCTGGAGACTTCTGAACGA | CCGTTCTTCACCGACTTCCT |
| *KRT7* | GGGAGCCGTGAATATCTCTGT | TGGAGAAGCTCAGGGCATTG |
